# Fc-dependent protective efficacy of non-inhibitory antibodies targeting influenza A virus neuraminidase is limited by epitope availability

**DOI:** 10.1101/2024.09.03.611041

**Authors:** Mirte N. Pascha, Marlies Ballegeer, Rien van Haperen, Annick C. Kooij, Danique M. van Miltenburg, Anthony A. Smits, Jelle G. Schipper, Hongrui Cui, Irina C. Albulescu, Berend Jan Bosch, Frank Grosveld, Frank J.M. van Kuppeveld, Xavier Saelens, Dubravka Drabek, Cornelis A.M. de Haan

## Abstract

Antibodies targeting hemagglutinin and neuraminidase (NA) are key components of the adaptive immune response against influenza A virus (IAV). However, antigenic drift allows the virus to escape inhibition by such antibodies. In this study, we aimed to isolate antibodies with cross-subtype reactivity against human H1N1 and H3N2 IAVs from transgenic mice bearing genes encoding the human immunoglobulin variable regions. We immunized these mice with recombinant N1 and N2 NA proteins, presenting them either as unconjugated soluble proteins or conjugated to self-assembling protein nanoparticles. This approach yielded a panel of NA-specific monoclonal antibodies (mAbs) with various levels of intra-and inter-subtype reactivity for N1 and N2 NA. Three of these mAbs, which collectively recognize two distinct epitopes, were cross-reactive against N1 and N2 NAs in ELISA, but did not inhibit NA enzymatic activity. Two of these mAbs, 21H8 and 45D9, were selected for further characterization. These recognized different epitopes and induced Fc-mediated effector functions to varying extents. Prophylactic administration of 21H8, but not 45D9, protected mice against challenge with H1N1 IAV, while neither mAb protected against a H3N2 challenge. The observed protective efficacy correlated with the mAbs’ capacity, or lack thereof, to bind membrane-associated full-length NA. The introduction of Fc silencing mutations in mAb 21H8 resulted in an inability to activate NK cells or mediate phagocytosis *in vitro* and significantly reduced protection *in vivo*, indicating that the protective efficacy of mAb 21H8 is Fc-dependent. However, mAb 21H8 expressed with reduced core fucosylation of its Fc N-glycan, which specifically enhanced NK cell activation *in vitro*, failed to improve protection against H1N1 challenge *in vivo*. Future work is needed to decipher in more detail the mechanism of Fc-mediated protection against influenza via NA-specific antibodies and to identify the optimal strategies for their enhancement.

## Introduction

Influenza A virus (IAV) is a highly contagious respiratory pathogen that causes seasonal epidemics and occasional pandemics, posing a significant threat to public health worldwide. Human IAV is notorious for its rapid antigenic evolution, resulting in the emergence of novel strains that can evade pre-existing immunity. Current inactivated vaccines mainly induce antibodies directed against immunodominant epitopes of the main surface glycoprotein hemagglutinin (HA), which are highly variable.^1^ Vaccine efficacy is therefore largely restricted to the strains included in the vaccine, resulting in a need for novel vaccines that induce a more broadly, cross-reactive immune response.^2,3^ The discovery of novel, broadly protective antibodies could aid in vaccine development and advance the development of antibody-based therapeutics.

Besides the attachment and fusion protein HA, neuraminidase (NA) plays a critical role in the viral life cycle.^4^ NA cleaves sialic acid residues from (decoy) receptors, facilitating the release of newly formed virus particles from infected cells and movement through the sialic acid-dense environment of the respiratory tract to infect new host cells. In line with this essential role in the virus life cycle, antibodies targeting NA can reduce disease severity and virus transmission.^5–9^ NA-targeting antibodies may block enzymatic activity and thereby interfere with viral release and spread, but their antiviral potential extends beyond direct inhibition.^10,11^ Antibodies with no or limited NA inhibiting (NAI) activity have been shown to rely on their ability to activate the immune system through Fc effector functions for protection.^10–12^

By binding to Fc gamma receptors (FcγR) on immune cells, such as natural killer (NK) cells and macrophages, antibodies can trigger antibody-dependent cellular cytotoxicity (ADCC) and antibody-dependent cellular phagocytosis (ADCP). ADCC leads to the destruction of infected cells, while ADCP involves the engulfment and clearance of viral particles or virus infected cells by phagocytic cells.^13,14^ In addition, antibody binding to an infected cell can trigger complement deposition, resulting in complement-dependent cytotoxicity (CDC).^15^ These processes also result in the release of antiviral cytokines that further enhance immune activation.^16^ Interaction with FcγRs is required for the *in vivo* protective efficacy of non-neutralizing antibodies against IAV, including antibodies targeting the HA stem and NA-specific antibodies.^10,11,17^

NA is increasingly recognized as an attractive target for improved influenza vaccines.^18^ Advancing our knowledge on the characteristics of protective cross-reactive monoclonal NA-targeting antibodies may inform vaccine design to promote the induction of protective antibodies against conserved epitopes. Few NA-targeting antibodies have been described that display cross-subtype specificity. All reported cross-subtype-reactive monoclonal antibodies (mAbs) target the highly conserved catalytic site and inhibit NA activity by blocking the interaction with sialic acid.^19–21^ These mAbs were isolated from patients infected with seasonal IAV or a healthy donor with unspecified exposure history. The most recently reported catalytic site-targeting antibody binds through sialic acid receptor mimicry and thereby displays exceptional breadth across IAV and IBV NAs.^21^ Antibodies binding elsewhere on NA can be strain-specific or cross-reactive within the subtype, but generally do not bind multiple NA subtypes.^12,22–28^

We previously demonstrated that immunization with NA conjugated to Mi3 self-assembling protein nanoparticles induced higher antibody titers than unconjugated NA and provided superior protection against a lethal viral challenge.^29^ In addition, differences in reactivity against heterologous NA proteins indicated that nanoparticle conjugation of NA induced an antibody response characterized by altered epitope targeting compared to unconjugated NA. In this study we aimed to identify NA-targeting antibodies with cross-subtype protective efficacy. To this end, we immunized Harbour H2L2 transgenic mice with recombinant N1 and N2 NAs administered as unconjugated protein or conjugated to nanoparticles. These mice carry the human heavy and light chain variable regions and the rat immunoglobulin constant regions. The human/rat chimeric antibodies that they produce can be engineered into fully human antibodies by replacing the rat constant regions with that of any human immunoglobulin isotype and subclass. Here, we isolated a panel of monoclonal antibodies (mAbs) that displayed varying levels of cross-reactivity against N1 and N2 recombinant proteins and engineered them into human IgG1 antibodies. While some mAbs displayed broad cross-subtype reactivity, their binding to recombinant NA proteins in enzyme-linked immunosorbent assays (ELISA) did not correlate with binding of NA in the native conformation or with protective efficacy. One mAb (21H8) protected mice against a challenge with H1N1, but not H3N2 IAV, while another mAb (45D9), targeting a different epitope, did not provide protection against either virus. We demonstrate that 21H8 lacks detectable NAI activity and relies on Fc-effector functions to mediate protection. However, enhancing ADCC by reducing Fc fucosylation did not improve its potency.

## Results

### Production of recombinant NAs and nanoparticles

To elicit and isolate novel mAbs against N1 and N2 NA and compare epitope targeting between different antigen preparations, we generated recombinant NA antigens N1 WI13 (derived from A/Wisconsin/09/2013) and N2 GE18 (derived from A/Germany/7830/2018), unconjugated and conjugated to nanoparticles (NA-NP). The recombinant tetrameric NAs were produced in a eukaryotic expression system using a modification of a previously described construct design consisting of the NA ectodomain preceded by a tetrabrachion tetramerization domain.^30^ The constructs also included a Strep tag for affinity purification and a SpyTag for nanoparticle conjugation (Fig. 1a), as described previously.^31^ Mi3-SpyCatcher^32^ and lumazine synthase (LS)-SpyCatcher self-assembling protein nanoparticles (NPs) were expressed in *E.coli* and affinity purified using a C-tag. Conjugation of the NA antigens to Mi3 or LS NPs occurred by spontaneous formation of an isopeptide bond between the SpyTag and SpyCatcher^33^ upon their coincubation (Fig. 1b, Fig. S1).

**Figure 1.**
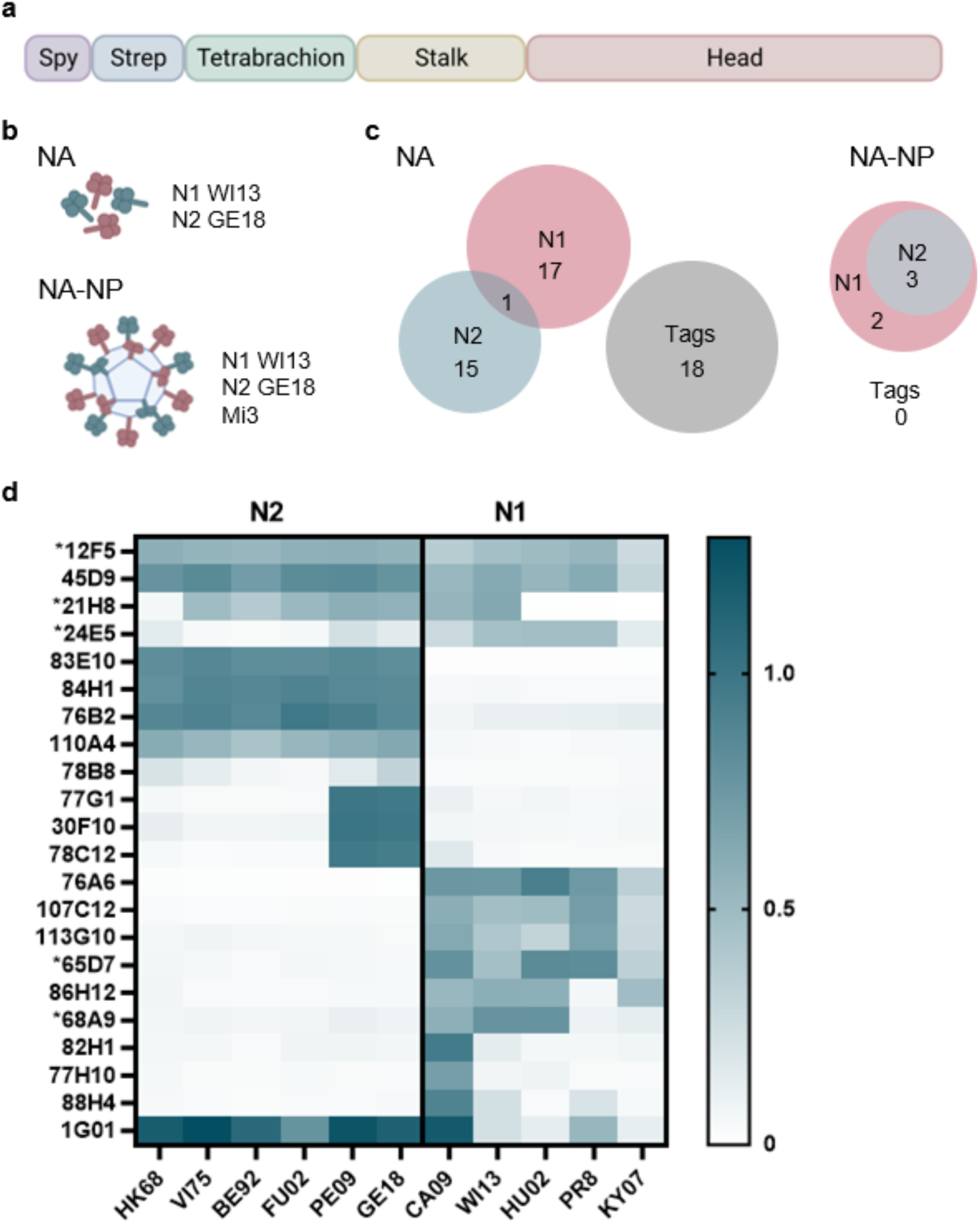
NA antigen design and antibody specificity. Recombinant tetrameric N1 and N2 NA antigens were produced and used to immunize H2L2 transgenic mice. (a) Schematic representation of the NA construct design. The constructs consist of the full NA ectodomain including the head and stalk, with an N-terminal tetrabrachion tetramerization domain, Twin-Strep-tag, and SpyTag. (b) Schematic of the immunogens. The mice received a mixture of equimolar amounts of N1 WI13 and N2 GE18 recombinant NAs, either unconjugated (NA) or conjugated to self-assembling protein nanoparticles (NA-NP). (c) Antigen specificity of hybridomas generated from immunized mice. Numbers indicate the amount of hybridomas positive for the indicated antigen in an ELISA assay. Tags refers to hybridoma supernatants binding the SpyTag, Strep-tag, or tetramerization domain. (d) Reactivity of purified antibodies (1 µg/ml) against N1 and N2 NAs of different strains measured in ELISA (OD at 450 nm, corrected for antigen coating as described in Materials and Methods, mean of 3 replicates). Strain identities are indicated in the Materials and Methods section. 1G01: control cross-reactive mAb [Stadlbauer]; * antibody isolated from mice immunized with NA-NP.

### Immunizations of H2L2 mice

We immunized Harbour H2L2 mice (6 per group) with a mixture of unconjugated N1 and N2 or with N1 and N2 co-conjugated to Mi3 or LS nanoparticles via subcutaneous immunization of 25 µg NA per mouse formulated with oil-in-water-based adjuvants. The mice received a total of six doses administered at regular intervals of two weeks. For mice that received NA-NP, we alternated between NA-Mi3 and NA-LS to minimize anti-carrier responses. NA-specific antibody titers were monitored during immunizations in enzyme-linked immunosorbent assays (ELISA). Analysis of serum samples taken after the fourth immunization demonstrated reactivity against the homologous NAs (N1 WI13 and N2 GE18; Fig. S2a and c), varying degrees of cross-reactivity against heterologous N1 and N2 proteins (N1 HU02 and N2 SI57; Fig. S2b and d), and low cross-subtype reactivity against N9 (Fig. S2e). High reactivity was also detected against the tetramerization domain and tags fused to the NA proteins, as demonstrated by binding to an unrelated RSV G recombinant protein containing the same domains as the NAs (Fig. S2f).

### Screening of NA-specific mAbs

After six immunizations of the H2L2 mice with NA or NA-NP antigens, B cells were isolated and fused with a myeloma cell line into hybridomas expressing monoclonal antibodies. We then screened nearly 12,000 hybridoma supernatants for binding to NA in an antigen-specific ELISA. In total, 51 hybridomas derived from mice immunized with unconjugated NA proteins demonstrated reactivity against N1 and/or N2. Of these, 18 also showed reactivity against the RSV G control protein, indicating that these antibodies interacted with the tags or tetramerization domain rather than NA. Of the 33 remaining NA-specific mAbs, one was cross-reactive against N1 and N2, 17 were specific for N1, and 15 were specific for N2. Immunizations with NA-NP yielded only five reactive hybridomas. Of these, three cross-reacted with N1 and N2 NA, two were specific for N1 NA, and none bound the recombinant RSV G control protein (Fig. 1c).

### Selection of cross-reactive mAbs

Next, we sequenced the genes encoding the IgG heavy and light chains of the mAbs specific for NA. Some overlap in sequences was found, resulting in a selection of 21 IgG mAbs with unique CDR region sequences. These selected mAbs were then purified from the supernatant for further profiling of their antigen specificity. Reactivity against a range of N1 and N2 NA proteins was determined by ELISA using the mAb 1G01^19^, a broadly neutralizing anti-NA mAb with cross-subtype specificity, as a control (Fig. 1d). Based on antigen specificity, the 21 mAbs could be roughly grouped into five categories; those with cross-subtype reactivity, broad reactivity within the N2 subtype, specific reactivity against N2 of recent H3N2 strains, broad reactivity within the N1 subtype, and specific reactivity against N1 of pandemic H1N1 strains. Two of the mAbs with cross-subtype specificity (12F5 and 45D9) also demonstrated broad reactivity within the N1 and N2 subtypes. 21H8 was broadly reactive within the N2 subtype, but only bound N1 NAs of new pandemic H1N1 origin. 24E5 was broadly reactive within the N1 subtype, but only demonstrated weak binding against a few N2 NAs (Fig. 1d; Fig. S3).

### Binding and inhibition characterization of the cross-reactive mAbs 12F5, 21H8, and 45D9

We moved forward with mAbs 12F5, 21H8, and 45D9 as these demonstrated the strongest binding to both N1 and N2 NAs. These mAbs were recombinantly expressed as human IgG1 isotypes for further characterization. The hIgG1 mAbs displayed high affinity binding to N1 WI13 in ELISA, with half-maximum binding values (EC_50_) of 100.7 ng/ml for 12F5, 94.6 ng/ml for 21H8, and 368.2 ng/ml for 45D9 (Fig 2a). For N2 GE18, the EC_50_ values were 47.6 ng/ml for 12F5, 134.4 ng/ml for 21H8, and 202.8 ng/ml for 45D9 (Fig. 2b). These findings confirmed our data collected with the rat IgG isotype mAbs (Fig. S4). We then assessed whether the mAbs were able to inhibit sialidase activity of NA in an enzyme-linked lectin assay (ELLA). While the 1G01 mAb effectively inhibits NA activity by binding directly to the catalytic site^19^, no NA inhibiting (NAI) activity of H1N1 or H3N2 was observed for 12F5, 21H8, or 45D9 in an enzyme-linked lectin assay (ELLA) (Fig. 2c-d; Fig. S4).

**Figure 2.**
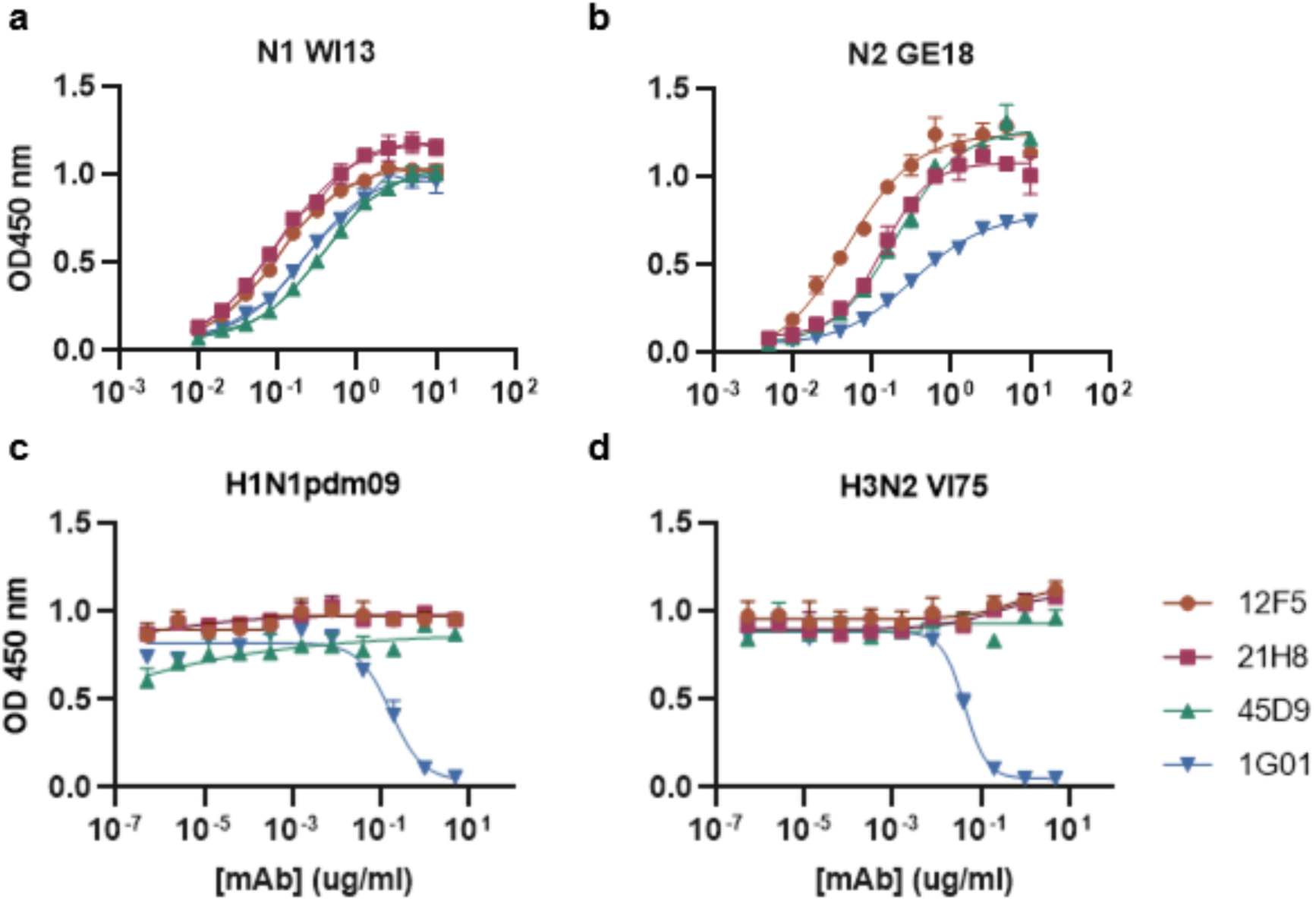
Binding-and functional characterization of cross-reactive hIgG1 mAbs. Recombinant purified human IgG1 mAbs (12F5, 21H8, 45D9, and positive control mAb 1G01) were further characterized. Binding reactivity was quantified in an ELISA against N1 WI13 (a) and N2 GE18 (b) (representative data of two independent experiments, mean ±SD of two replicates). Inhibition of NA activity was determined in an ELLA with H1N1pdm09 (c) and H3N2 VI75 (d) (mean ±SD of three replicates).

### MAbs 12F5, 21H8, and 45D9 bind linear epitopes

The lack of NAI activity by the cross-reactive mAbs is suggestive of binding to an epitope distant from the catalytic site. To elucidate their epitopes, we first assessed the ability of the mAbs to bind denatured NA. 12F5, 21H8, and 45D9 all bound N1 and N2 NAs that were denatured by heat treatment in the presence of sodium dodecyl sulfate and 2-mercaptoethanol (Fig. 3a-b), suggesting that these mAbs bind linear sequences in the NAs. In contrast, the 1G01 mAb, which is known to bind a conformational epitope^19^, did not display any binding against denatured NA. As expected, an antibody recognizing the linear strep tag displayed high reactivity against denatured NA.

**Figure 3.**
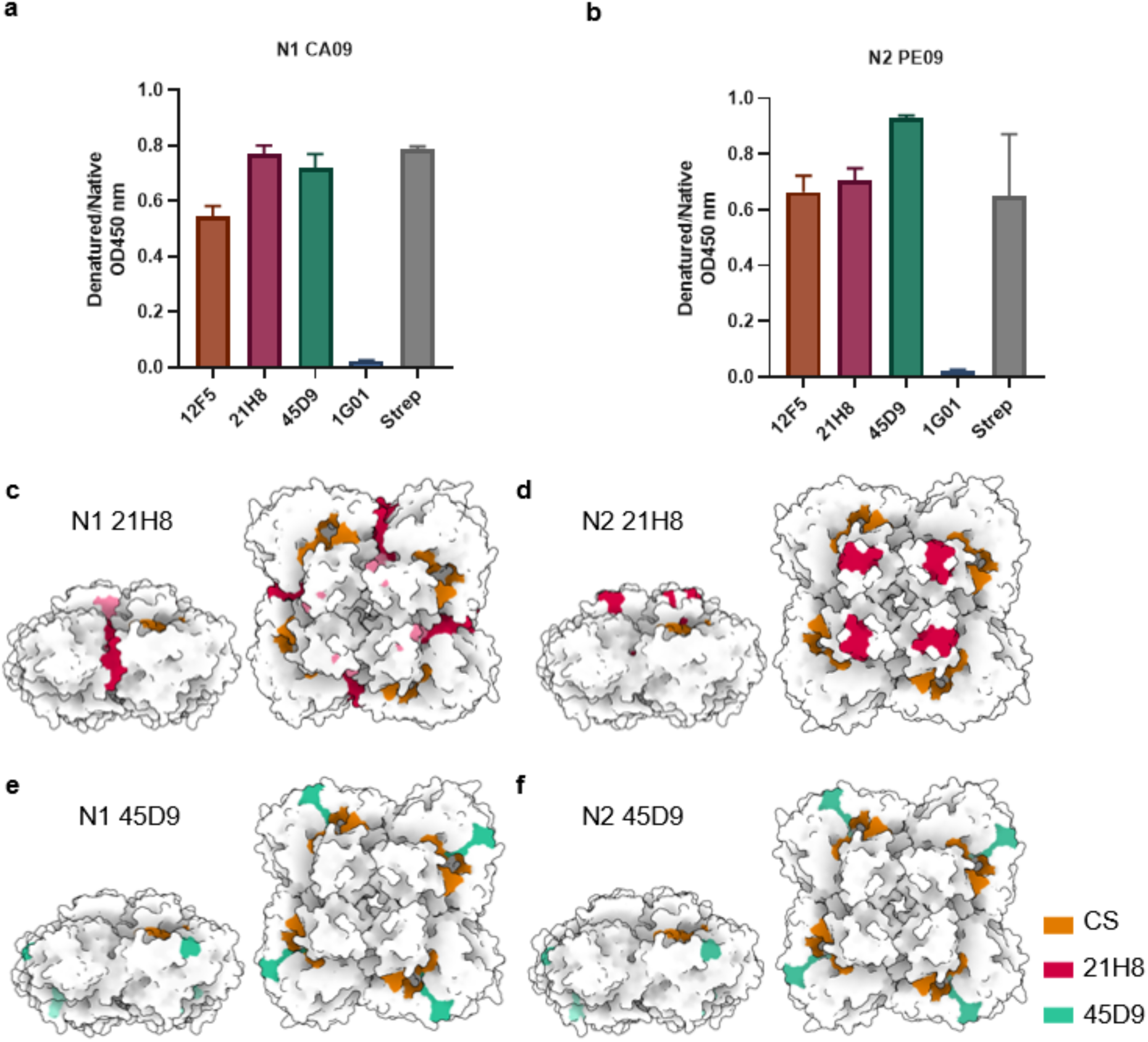
Cross-reactive mAbs bind linear epitopes on N1 and N2. (a and b) Binding of mAbs to denatured recombinant NA proteins (a, N1 CA09; b, N2 PE09) was analyzed relative to their binding to the proteins in their native conformation in ELISA to determine whether mAbs bound conformational or linear epitopes (mean ±SD of three replicates). 1G01: NA mAb binding a conformational epitope; Strep: mAb binding the linear sequence of the StrepTag. (c to f) Residues corresponding with pepides recognized by mAbs 21H8 and 45D9 in peptide arrays (Fig. S5 and S6) are indicated on the structures of N1 and N2. Catalytic site (CS) indicated in orange, epitopes in pink (21H8) and green (45D9). When the antibodies recognized two peptides, the overlapping sequence is indicated with a darker shade and the remaining parts of the sequences with a lighter shade. Sequences of the peptides are indicated in Fig. S7. N1 structure: PDB 3NSS [Li Nat Struct Mol Biol 2010]. N2 structure: PDB 4GZP [Zhu J Virol 2012].

Since all three mAbs bound a linear amino acid sequence within the NAs, we were able to map the epitopes with tiled peptide arrays. Binding of the mAbs to 13-to 15-meric peptides spanning the sequences of N1 (A/New York/18/2009) and N2 (A/Perth/16/2009) with 11 amino acids overlap was assessed by ELISA. 21H8 bound to peptides corresponding to amino acids 446-460 and 450-464 in N1. This mAb was not reactive against the corresponding peptides in N2, but instead bound a peptide corresponding to an adjacent region (433-447) with no shared sequence identity except for two overlapping amino acids (Fig. S5a and S6a). The peptide sequence of N1 corresponds to the lateral interface between the NA protomers and extends towards the carboxy-terminus (C-terminus) on top of the protein (Fig. 3c). The peptide sequence of N2 corresponds to an internal ß-strand and a loop at the top of the protein adjacent to the C-terminus (Fig. 3d). 45D9 bound peptides corresponding to amino acids 288-302 and 292-306 (N1), and 289-303 and 293-306 (N2) (Fig. S5c and S6c). These regions map to two beta strands inside the NA head and a surface loop adjacent to the catalytic site (Fig. 3e-f). 12F5 was found to bind a similar region as 45D9 in N1 (amino acids 288-302) and N2 (amino acids 293-306) (Fig. S5b and S6b). The identified peptide sequences are highly conserved within the subtypes and, for the most part of the sequences recognized by 12F5 and 45D9, also between N1 and N2 (Fig. S7). As 45D9 and 12F5 bind to similar epitopes, we continued the characterization with 45D9 and 21H8.

### MAbs 21H8 and 45D9 induce Fc-mediated effector functions

In the absence of direct neutralizing capacity, anti-NA mAbs may protect through Fc-mediated immune activation. MAbs 21H8 and 45D9 were both able to activate NK cells *in vitro* when bound to N1 or N2 recombinant proteins adsorbed to ELISA plates (Fig. 4a and c). NK cell activation against N2 VI75 was higher for 45D9 than for 21H8, whereas 21H8 induced higher activation against N1 CA09, consistent with their respective binding strengths against these NAs in ELISA (Fig. 4b and d). Binding of 21H8 to N1 NA immobilized on beads also effectively induced phagocytosis by THP-1 cells and, in higher concentrations, complement deposition (Fig. 4e-f). 45D9 demonstrated limited binding to N1 conjugated to beads (Fig. 4g) and, probably as a result thereof, did not induce phagocytosis or complement activation in these assays as efficiently as 21H8 (Fig. 4e-f). Coupling the NA proteins to the neutravidin beads required their biotinylation. To this end, we incorporated a biotin acceptor peptide (BAP) in the NA constructs and co-expressed these with biotin ligase (BirA), resulting in the in situ site-specific biotinylation of the NAs. This biotinylation process impacted the expression levels of the NA proteins considerably. Consequently, we were unable to recover adequate amounts of the N2 NAs to conduct the assays with.

**Figure 4.**
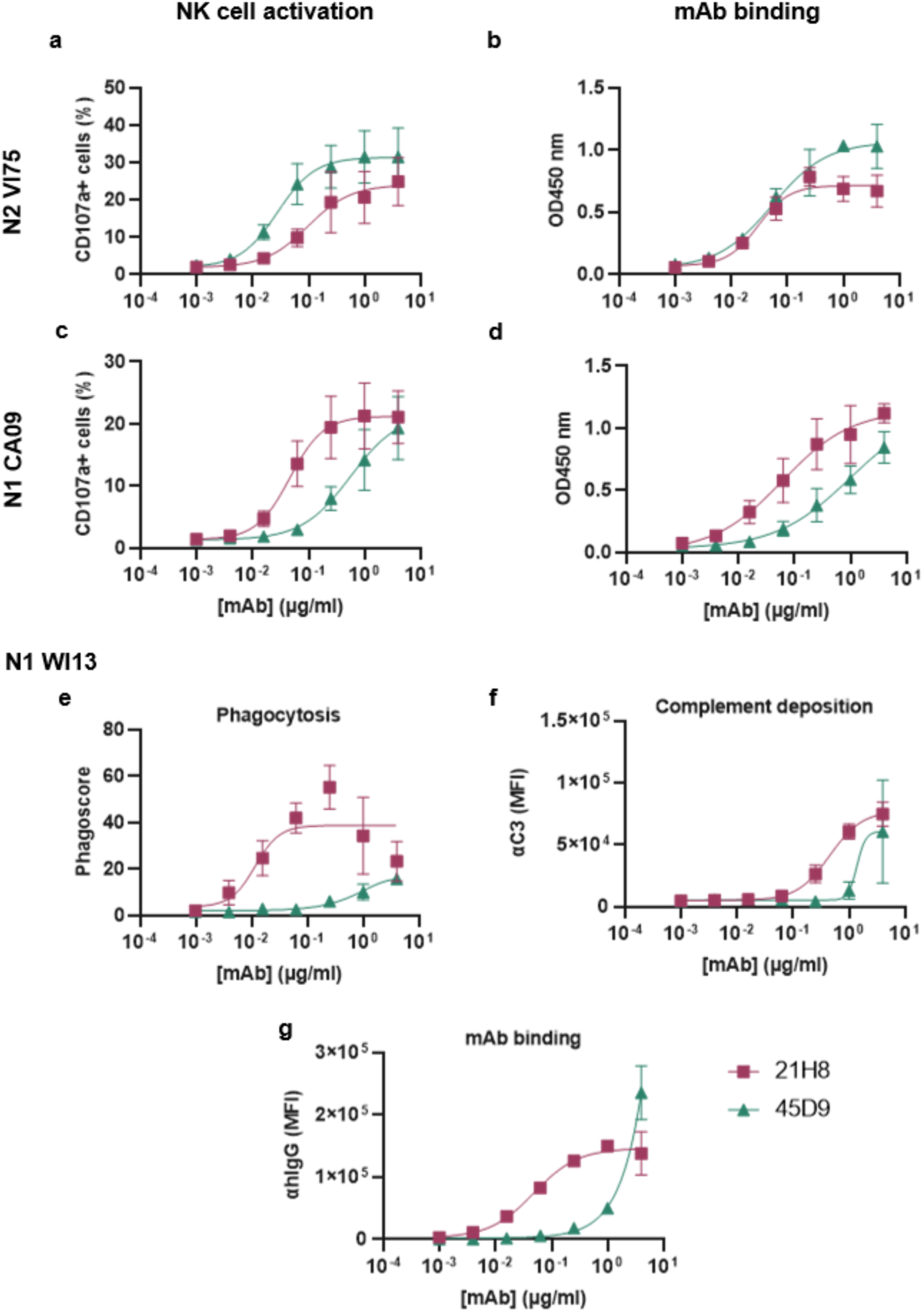
21H8 and 45D9 induce Fc effector functions *in vitro*. (a-d) The ability of the mAbs to induce activation of NK92 cells was determined following the incubation of the cells in plates coated with N1 CA09 or N2 VI75 in the absence or presence of the mAbs. The amount of activated cells was measured as CD107a^+^ cells in flow cytometry and graphed as a percentage of the total NK cells (a and c). Binding of the mAbs to the NAs was measured in ELISA (b and d). (e to g) The induction of phagocytosis and complement deposition by the mAbs was determined in flow cytometry-based assays using microspheres. (e) Phagocytosis of microsphere-based immune complexes containing N1 WI13 and the mAbs by THP-1 monocytes was measured by flow cytometry. The phagoscores were calculated based on the percentage of THP-1 cells positive for the uptake of microspheres and the gMFI of that population as detailed in the Materials and Methods section. (f) Deposition of complement protein C3 on the microsphere-based immune complexes was measured by flow cytometry. (g) mAb binding to the N1-coated microspheres measured by flow cytometry. Data shown represent the mean ±SD of in total four replicates from two independent experiments combined.

### 21H8, but not 45D9, protects from IAV infection

To assess whether the mAbs could protect *in vivo*, we prophylactically treated mice with 21H8 or 45D9 before challenge with H1N1 or H3N2 IAV. Mice (n = 6 per group) were treated intranasally with 50 µg of the anti-NA hIgG1 21H8 or 45D9, a positive control mAb (N1-C4 NAI mAb^22^) for the H1N1 challenge virus, or a negative control irrelevant hIgG1 one day prior to virus challenge. hIgG1 binds mouse Fc receptors with affinities similar to the mouse isotypes^34^ and effectively induces Fc effector functions in murine effector cells^35^, indicating that hIgG1 mAbs can be used in mice for the detection of Fc-mediated protection.

mAb 21H8 effectively protected mice from H1N1 challenge. Mice that received this antibody prior to infection experienced limited weight loss and survived the challenge (Fig. 5a-b). However, weight loss was somewhat higher than in the mice treated with the positive control NAI mAb (Fig. 5a; table S2). In contrast, 21H8 failed to protect mice from H3N2 infection. These mice all lost weight and died or reached the ethical endpoint at a rate similar to those that received the negative control mAb (Fig. 5c-d). MAb 45D9 provided very limited, if any, protection against either IAV challenge. Most mice that received 45D9 lost weight similar to the negative control treated mice, although a few mice experienced reduced weight loss (Fig. 5a and c; table S2). In addition, three out of six mice treated prophylactically with 45D9 survived in both the H1N1 and H3N2 experiments, compared to one of the negative control-treated mice infected with H1N1 and two in the negative control group infected with H3N2 (Fig. 5b and d).

**Figure 5.**
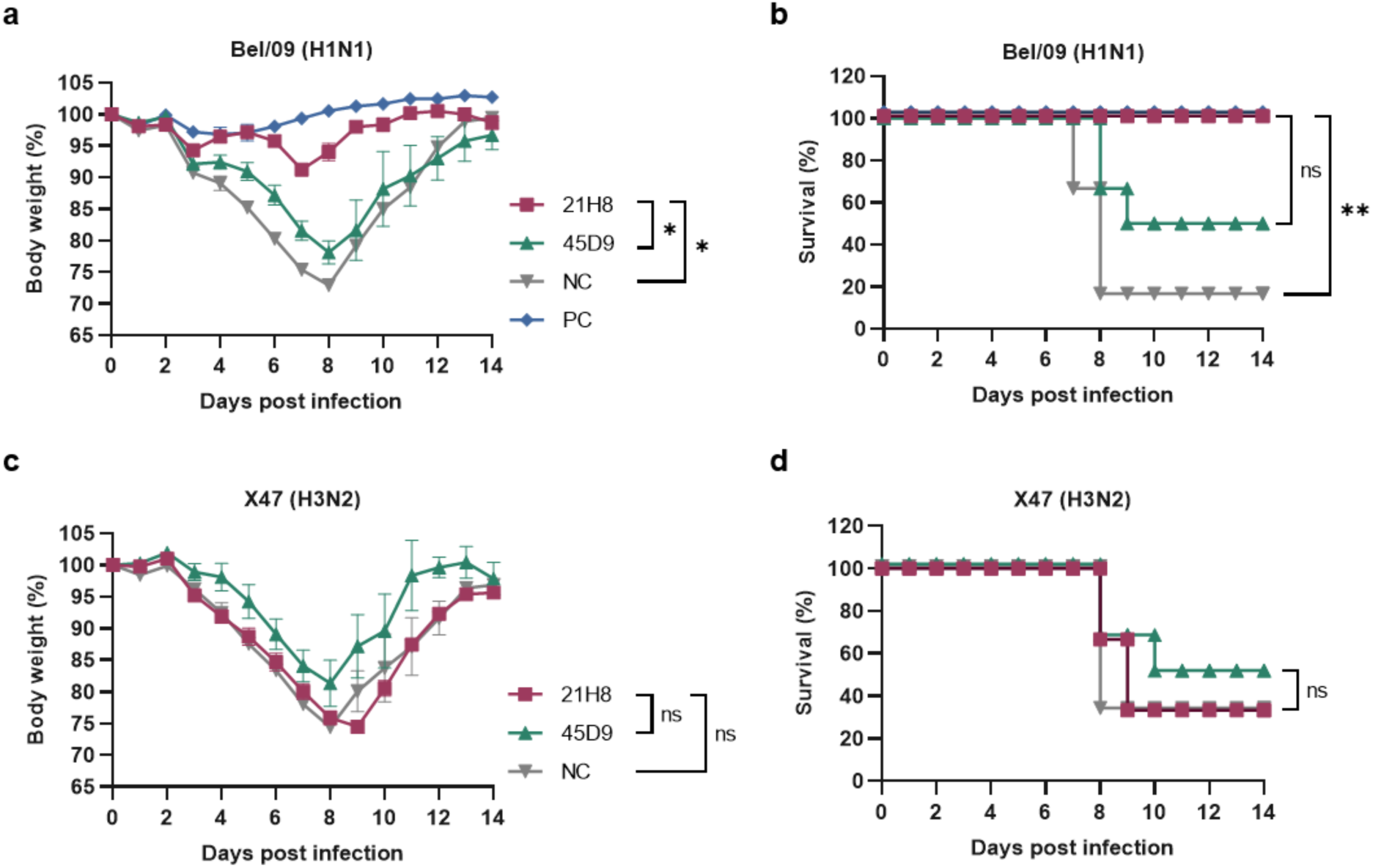
*In vivo* prophylactic effect of 21H8 and 45D9 against IAV challenge in mice. Challenge with 2LD_50_ of mouse-adapted H1N1 (Bel/09) and H3N2 (X47) virus after prophylactic administration of 50 µg 21H8, 45D9, isotype control mAb (anti-RSV hIgG1; NC), or positive control mAb (N1-C4; PC) one day prior. BALB/c mice (6 per group) were monitored daily for body weight (a and c; weight relative to baseline, displayed as mean ± SEM) and survival (b and d; expressed as percentage of the group) for 14 dpi. Body weight data were analyzed using the method of residual maximum likelihood (REML) for repeated measurements (ns, not significant; *, F pr. < 0.001), and survival data were analysed using the Mantel-Cox log-rank test (ns, not significant; *, p < 0.05; **, p < 0.002; ***, p < 0.0002).

Collectively, these results demonstrate protective efficacy *in vivo* only for mAb 21H8 against H1N1 IAV challenge. 21H8 most likely protects against H1N1 through Fc effector functions but was unable to protect against H3N2 infection. Protective efficacy of 45D9 was very limited against both H1N1 and H3N2, despite its strong NK cell-activating effect particularly when bound to immobilized N2 NA *in vitro*.

### Protection by 21H8 and 45D9 mAbs correlates with binding to NA on virus or to infected cells

Further investigation into the lack of correlation between *in vitro* Fc effector functions and *in vivo* protection of our mAbs led us to examine their binding to NA in a more biologically relevant context. Whereas both 21H8 and 45D9 demonstrated strong binding by ELISA (Fig. 2a-b) to recombinant NA proteins, which have been confirmed to be well-folded and enzymatically active^29,36^, binding against NA on IAV particles was much lower (Fig. 6a-b). Consistent with the challenge results, 21H8 demonstrated good binding to H1N1 but not to H3N2. Binding of 45D9 was very low against H1N1 and not detectable against H3N2.

**Figure 6.**
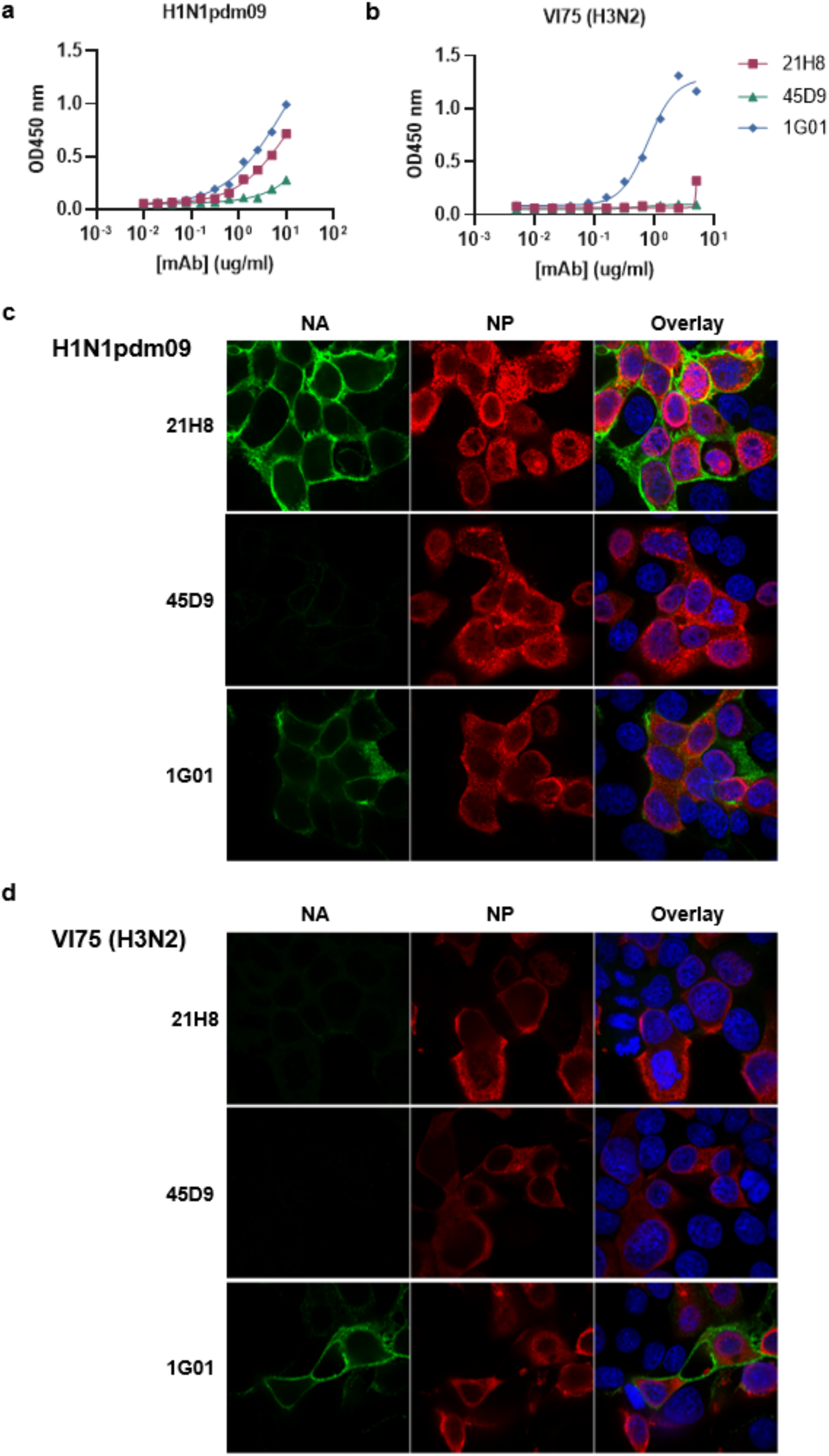
Binding of 21H8 and 45D9 to full-length NA in membranes of viruses and infected cells. (a and b) Binding of the mAbs to NA of H1N1pdm09 (a) and H3N2 VI75 (b) determined in ELISA with whole viruses adsorbed to the plates (representative results of two independent experiments are shown). (c and d) Immunofluorescence analysis of MDCK cells infected with H1N1pdm09 (c) and H3N2 VI75 (d). Surface staining with the NA-targeting mAbs (green) and intracellular staining with a mAb targeting the nucleoprotein (NP; red). Nuclei are labelled with dapi (blue). Representative pictures are shown.

These observations were confirmed for NA expressed on infected MDCKII cells. 21H8 and 1G01, but not 45D9, were able to bind NA on the surface of H1N1 infected cells (Fig. 6c). Some low reactivity with 45D9 was detected in permeabilized H1N1 infected cells, indicating that this mAb can bind the NA only intracellularly (Fig. S8). 1G01 was able to bind NA on the surface of H3N2 infected cells, indicating the expression and correct folding of the NA. However, no reactivity of our mAbs 21H8 and 45D9 was detected against cells infected with this virus (Fig. 6d and Fig. S8).

### Fc engineering to modulate Fc effector functions of 21H8

To increase the potency of 21H8 and improve our understanding of the role of Fc effector functions in protection against IAV, we modulated the Fc domain of 21H8 and tested different variants in the H1N1 challenge model. First, we introduced the silencing mutations LALA-PG (L234A, L235A, and P329G) to confirm that 21H8 protects against H1N1 infection solely through Fc effector functions. The LALA-PG mutations eliminate complement binding and interaction with Fcγ receptors of hIgG1, resulting in a complete loss of Fc effector functions.^37^ Second, to determine whether protective efficacy could be improved by enhancing ADCC, we modified the glycosylation of 21H8. The absence of a core fucose on the N-linked glycan at position N297 in the interface between the Fc protomers of hIgG1 enhances affinity for activating FcγRs associated with ADCC activity (FcγRIIIa in humans).^38^ We produced 21H8 with low fucose content (21H8ΔF) by co-expression of GDP-6-deoxy-D-lyxo-4-hexulose reductase (RMD)^39^ in cells expressing the mAb. Reduced fucose content was confirmed by ELLA using *Aleuria aurantia* lectin (AAL) (Fig S9).

The Fc modifications of 21H8 hIgG1 had the expected effects on modulating NK cell activation *in vitro* without affecting the binding affinity of the mAbs against recombinant N1 protein (Fig. 7a). 21H8-LALA-PG was unable to activate NK cells, whereas the number of activated (CD107a^+^) NK cells more than doubled with 21H8ΔF compared to the wild type 21H8 with high fucose content (Fig. 7b). The enhancing effect of the modification was even stronger for NK cell activation against the virus particles, to which the mAbs bound only to low levels probably due to low NA content in these preparations (Fig. 7c and d). 21H8ΔF induced phagocytosis to equal levels as the unmodified mAb, while phagocytosis was absent with 21H8-LALA-PG (Fig. S10).

**Figure 7.**
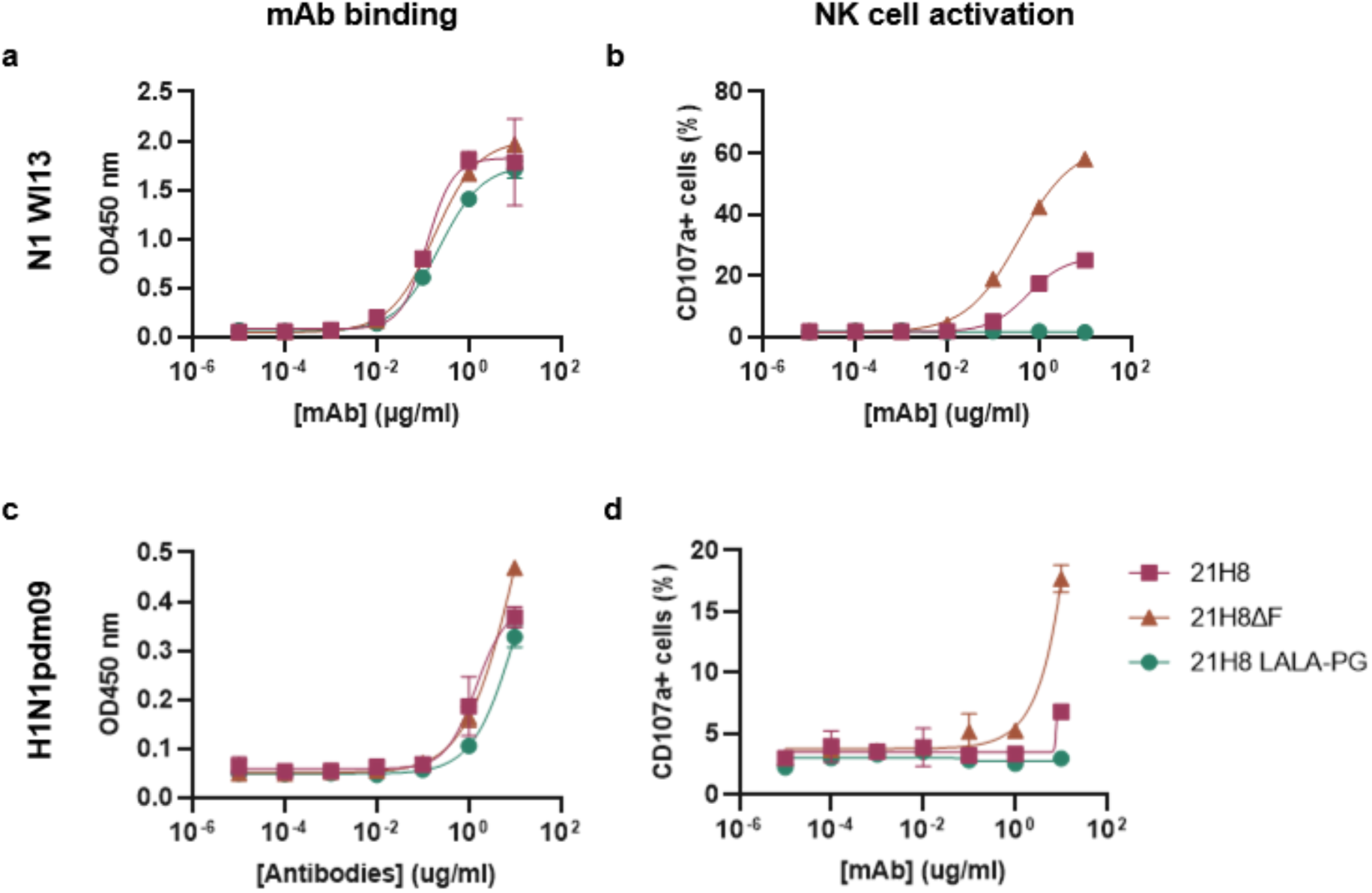
Fc modifications modulate NK cell activation by 21H8. 21H8 produced with reduced fucosylation (21H8ΔF) for enhanced FcγR engagement or mutated to abolish FcγR engagement (21H8 LALA-PG). Binding of the 21H8 variants against recombinant soluble N1 WI13 (a) and H1N1pdm09 virus (c) was quantified in ELISA. (b and d) The ability of the modified antibodies to activate NK cells upon binding to recombinant soluble N1 WI13 (b) or H1N1pmd09 virus (d) was quantified in flow cytometry as the percentage of CD107a^+^ NK cells. Representative data of two independent experiments are shown (mean ±SD of two replicate measurements).

After confirming the functional effects of the Fc modifications *in vitro*, we tested the prophylactic efficacy of 21H8ΔF and 21H8-LALA-PG relative to the wild-type, high fucose 21H8 mAb in the H1N1 challenge model. For this challenge experiment, we used a lower dose of the mAbs (10 µg instead of 50 µg in the experiment in Fig. 5) to discern potential positive effects of reduced fucosylation. The protective efficacy against challenge with H1N1 Bel/09 at either two-or four times the median lethal dose (2LD_50_ or 4LD_50_) was reduced significantly for 21H8-LALA-PG (Fig. 8, Table S3), suggesting that 21H8 indeed largely depends on Fc-mediated effector functions to mediate protection. However, the LALA-PG mutant retained a low level of protective activity against weight loss, which was significant only in the 2LD50 treated mice. While with both challenge doses all six mice that received the negative control IgG died after six to nine days, one or two mice that received the 21H8-LALA-PG mAb survived the challenge. Mice that received the unmodified 21H8 mAb experienced substantial weight loss, but most survived the challenge except for one mouse that died on day 10 post-challenge with 4LD_50_ Bel/09. Despite its enhanced ability to activate NK cells *in vitro*, 21H8ΔF did not display enhanced protection *in vivo* compared to the unmodified 21H8. Weight loss in mice of both groups was similar, although in the 2LD_50_ Bel/09 challenge dose setting, the mice treated with 21H8ΔF made a faster recovery when compared to the wild-type antibody. However, more mice died in the groups that received 21H8ΔF, one in the group challenged with 2LD_50_ and two in the group challenged with 4LD_50_ Bel/09 (Fig. 8).

**Figure 8.**
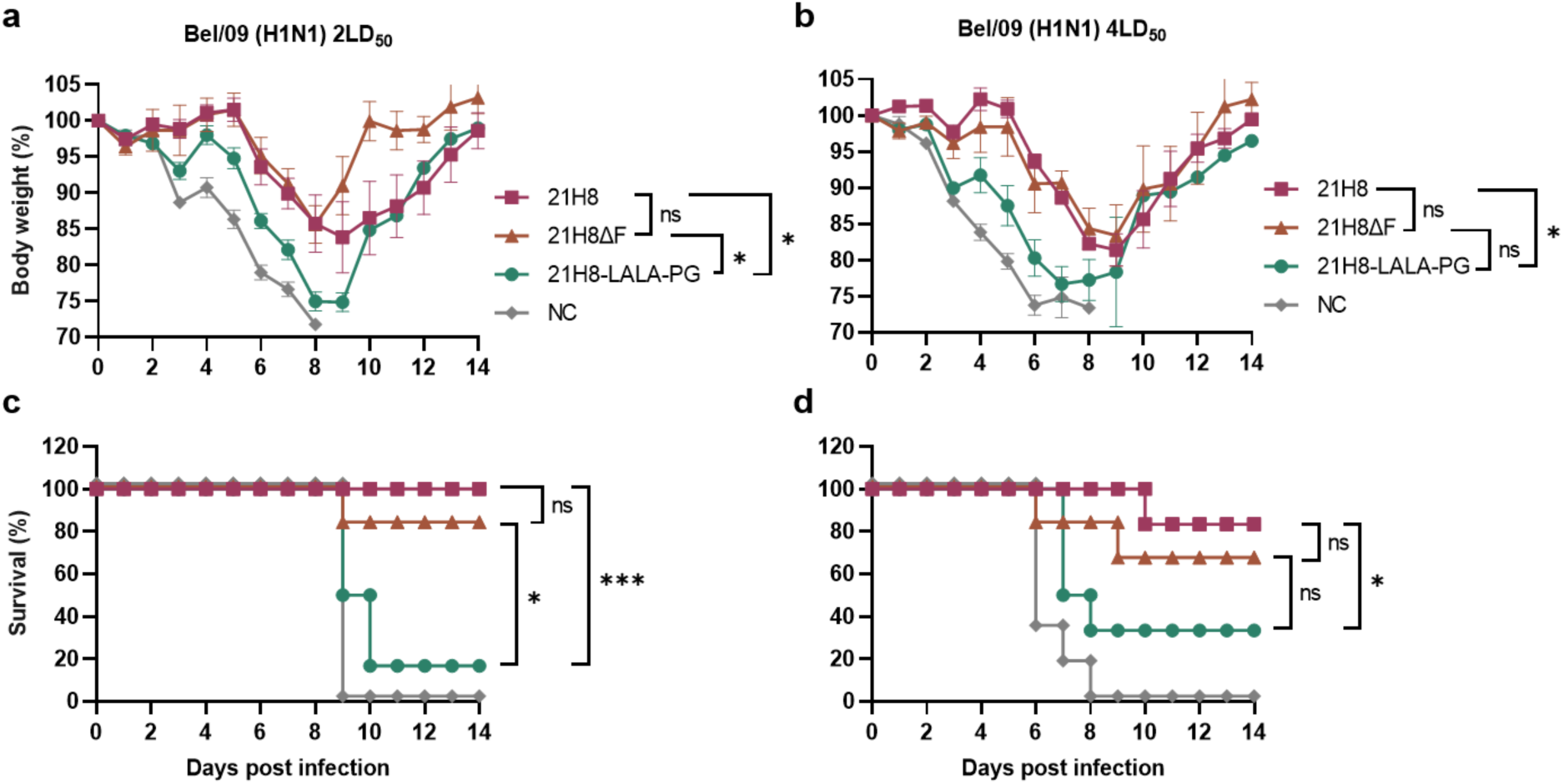
*In vivo* prophylactic effect of Fc modified 21H8 against H1N1 challenge in mice. Challenge with mouse-adapted H1N1 (Bel/09) virus at 2LD_50_ or 4LD_50_ after prophylactic administration of 10 µg 21H8, 21H8ΔF, 21H8-LALA-PG, or isotype control IgG (anti-RSV hIgG1; NC) one day prior to infection. BALB/c mice (6 per group) were monitored daily for body weight (a and b; weight relative to baseline, displayed as mean ± SEM) and survival (c and d; expressed as percentage of the group) for 14 dpi. Body weight data were analyzed using the method of residual maximum likelihood (REML) for repeated measurements (ns, not significant; *, F pr. < 0.001), and survival data were analysed using the Mantel-Cox log-rank test (ns, not significant; *, p < 0.05; ***, p < 0.0002).

## Discussion

Antibodies, including those targeting NA, are an essential component of immunity against IAV. The discovery of novel broadly protective antibodies may give rise to new antibody-based therapeutics. In addition, based on these antibodies novel conserved antigenic sites may be identified that can aid in the development of broadly reactive vaccine immunogens. Here, we immunized H2L2 transgenic mice with either unconjugated or nanoparticle-conjugated recombinant N1 and N2 NAs to elucidate putative differences in epitope targeting and to discover new antibodies that can recognize both NA subtypes. We identified several antibodies with varying breadth across different N1 and N2 strains, including four that display broad cross-subtype reactivity in ELISA. However, these cross-reactive antibodies appear to preferentially target occluded epitopes, resulting in reduced reactivity with membrane-bound NA, with the exception of mAb 21H8 binding to N1 NA. As a result, only 21H8 was able to protect mice against infection with H1N1, but not H3N2. This protection is mediated through Fc effector functions. We observed that reducing fucosylation of the glycan side chain in the Fc domain of 21H8 greatly enhanced NK cell activation *in vitro*. However, this modification did not improve protection *in vivo*.

We previously demonstrated that NA-Mi3 NPs induce higher NA-specific antibody titers than unconjugated NA antigens in Balb/c mice.^29^ While the overall antibody titers were generally lower in the Harbour H2L2 mice and the difference between NA-NP and unconjugated NA antigens was not significant, we did observe differences in antigen specificity which suggests an altered epitope targeting due to NP conjugation. Specifically, NA-NP induced more antibodies that displayed cross-subtype reactivity, whereas unconjugated NA primarily induced subtype-specific antibodies and antibodies targeting tags on the recombinant NA proteins. Nevertheless, these findings need to be interpreted with caution, as our analysis of antigen specificity was limited to mAbs expressed by hybridomas, which represents the polyclonal response incompletely. A comprehensive characterization of the antibody response through polyclonal epitope mapping using for example electron microscopy^40^ or hydrogen-deuterium exchange mass spectrometry^41^ could provide a more in-depth understanding of the changes in epitope targeting resulting from nanoparticle presentation.

The isolated mAbs 12F5 and 45D9 bound a linear epitope that is largely conserved between N1 and N2 NAs, resulting in cross-subtype reactivity against N1 and N2 NAs from various strains in ELISA. However, this binding was reduced considerably to NA proteins that were immobilized on a surface in a more defined orientation using a C-terminal peptide tag or biotinylation, as well as to full-length NA on IAV particles or infected cells. Our recombinant NA proteins adopt a closed tetrameric conformation resembling that of full-length membrane-bound NA and display high enzymatic activity^29^, suggesting that they retain their structural integrity in solution. Yet, the non-specific adsorption of the recombinant NA proteins onto the assay plates in ELISA may have compromised their conformation, leading to the exposure of otherwise occluded epitopes. This could have led to the identification of 21H8 and 45D9 as cross-subtype reactive mAbs, whereas their reactivity seems more restricted when NA is displayed in the native conformation. While 45D9 showed minimal binding to surface-expressed NA, it did exhibit some reactivity to intracellular NA. This might be attributed to the intracellular proteins being in the process of folding. Consistent with the observed lack of binding to surface-expressed NA, 45D9 was unable to protect against IAV challenge in mice.

MAb 21H8 also displayed cross-subtype reactivity against N1 and N2, but bound to different linear epitopes on N1 and N2 with very limited shared sequence identity. Since the N1 and N2 reactive sequences are adjacent to each other on the quaternary NA structure, this may indicate a larger antibody footprint with a different binding mode against each subtype. Similar to the epitopes of 12F5 and 45D9, the N2 epitope for 21H8 was unavailable for binding in the full-length NA, possibly due to it also being an occluded epitope. On N1, 21H8 binds 2 peptides located closer to the C-terminus when compared to N2. The overlapping peptide sequence shows a three amino acid overlap with the epitope identified for mouse mAb N1-7D3^22^, with the N1-7D3 mAb binding closer to the highly conserved C-terminus of N1 NA. Whereas N1-7D3 is broadly cross-reactive within the N1 subtype, including NA of pre-2009 H1N1 and avian H5N1, 21H8 binds only to NAs of post-2009 human H1N1 viruses.

The protective efficacy of 21H8 against a challenge with H1N1 in mice was largely dependent on the engagement of FcγRs, as demonstrated by the greatly reduced protection of the antibody variant harboring the LALA-PG Fc-silencing mutations. In agreement with these observations, the protective efficacy of the previously described mAb N1-7D3 that targets a partially overlapping epitope was dependent on the isotype. N1-7D3 is a mouse IgG1 antibody, which is a subtype known for weak induction of Fc effector functions, and therefore did not protect against IAV infection^22^, whereas a hIgG1 chimeric version of N1-7D3 did protect^11^. Similarly, the mAb 3C05, which showed broad reactivity within the N1 subtype and low neutralizing activity *in vitro*, was also dependent on FcR engagement for protection against H1N1.^10^

Since most activating FcγRs bind IgG with low affinity, the induction of Fc effector functions depends on multivalent interactions stemming from the clustering of IgG molecules on the surface of the virus or infected cells.^42^ The relatively low abundance of NA on these surfaces has been proposed to make NA-targeting antibodies less efficient inducers of Fc effector functions than HA-targeting antibodies.^43^ Aiming to augment the protective efficacy of 21H8, we reduced fucosylation of the N-linked glycan in the Fc domain. hIgG1 lacking the core fucose binds to human FcγRIIIa with higher affinity and thus more potently induces ADCC.^38^ The effects of the modification on other Fc effector functions like ADCP remain somewhat ambiguous but seem to be limited.^44^ Utilizing a human NK cell line, we confirmed the enhanced NK cell activation by the modified 21H8 mAb (21H8ΔF). Although the mouse orthologs of FcγRIIIa differ considerably from the human receptor, the enhanced stimulation of ADCC by afucosylated human IgG1 has been observed in different species, including mice.^45–47^

The reduced fucosylation of mAb 21H8 did not improve its potency against H1N1 challenge in mice. These findings align with the results of a study by Bournazos *et al.*, which described the effects of several different Fc modifications on the protective efficacy of HA-and NA-targeting antibodies.^48^ Enhancement of FcγRIIa binding augmented the protective efficacy in mice expressing human FcγRs, while modifications promoting binding to FcγRIIIa, including afucosylation, were ineffective. Collectively, our results and those from Bournazos and colleagues suggest that ADCC plays a minor role in antibody-mediated protection against influenza virus compared to other Fc effector functions. This interpretation aligns with observations from He *et al.*, which point towards a pivotal role of alveolar macrophages for antibody-mediated protection against influenza virus in mice, downplaying the role of NK cells.^49^ Nevertheless, in older human adults, NK cell activation and influenza-specific afucosylated antibodies have been demonstrated to correlate with enhanced protection.^50^ More studies are needed to further explore the importance of Fc effector functions in protection against IAV and of strategies to enhance the protective efficacy of non-neutralizing mAbs.

## Acknowledgements

The authors would like to thank Rory de Vries and Nella Nieuwkoop from Erasmus Medical Center, Rotterdam, and Puck van Kasteren, Anke Lakerveld, and Anne Gelderloos from the Rijksinstituut voor Volksgezondheid en Milieu (RIVM), Bilthoven for generous exchange of materials and technical advice. The authors are also grateful for the support of Richard Wubbolts from the Center for Cell Imaging (CCI) of the Faculty of Veterinary Medicine at Utrecht University. This study was done within the framework of the research program of the Netherlands Centre for One Health (www.ncoh.nl) and co-funded by the PPP Allowance made available by Health∼Holland (Grant no. LSHM19136), Top Sector Life Sciences & Health, to stimulate public-private partnerships. The following reagents were obtained through BEI Resources, NIAID, NIH: influenza virus peptide arrays A/New York/18/2009 (H1N1) (NR-19249) and A/Perth/16/2009 (H3N2) (NR-19267). M.B., H.C., and C.A.M.d.H. were supported by the ENDFLU project that received funding from the European Union’s Horizon 2020 research and innovation program under grant agreement No. 874650.

## Materials and Methods

### NA gene construct design

Human codon-optimized cDNA (Genscript, USA) encoding the NA ectodomains of A/Wisconsin/09/2013 (H1N1) (GenBank accession no. AGV29183.1; referred to as N1 WI13), A/California/04/2009 (H1N1) (GenBank: ACP41107.1; N1 CA09), A/Hunan/795/2002 (H5N1) (GenBank: BAM85820.1; N1 HU02), A/Puerto Rico/8/34 (H1N1) (GenBank: AAM75160.1; N1 PR8), A/Kentucky/UR06-0258/2007 (H1N1) (GenBank: CY028165.1; N1 KY07), A/Germany/7830/2018 (H3N2) (GenBank: QBH71200.1; N2 GE18), A/Netherlands/063/2011(H3N2) (GenBank: AFH00706.1; N2 PE09), A/Netherlands/213/2003(H3N2) (GenBank: AFN11848.1; N2 FU02), A/Netherlands/179/1993(H3N2) (GenBank: AFG72375.1; N2 BE92), A/Bilthoven/1761/1976(H3N2) (GenBank: AFG99020.1; N2 VI75), A/Hong Kong/1968 (H3N2) (GenBank: ABQ97206.1; N2 HK68) were cloned into pFRT expression plasmids (Thermo Fisher Scientific) as previously described.^30,51^ The NA ectodomain sequences were preceded by sequences encoding the signal peptide derived from Gaussia luciferase, a Twin-Strep-tag for affinity purification (IBA), and the Tetrabrachion tetramerization domain (similarly as described previously^30,51^). A RSV G ectodomain-encoding sequence^52^ was cloned in the same pFRT vector. The plasmids encoding N1 WI13 and N2 GE18 additionally contained the sequence encoding the SpyTag preceding the Strep tag sequence (as recently described^29^).

### Protein expression and purification

Proteins were expressed in FreeStyle 293-F cells (Gibco) that were maintained in Freestyle 293 Expression Medium (Gibco) at 37°C with 8% CO_2_ with shaking at 130 rpm. Plasmids encoding the proteins were transfected into the cells using polyethyleneimine (PEI) in a 1:3 ratio (µg DNA:µg PEI). After 24 hours Peptone Primatone RL (Sigma-Aldrich) and valproic acid (Sigma) were added to final concentrations of 0.5% and 2,25 mM, respectively. Cells were then incubated for a further 3-4 days until viability fell below 80%. Cell supernatant was harvested by centrifugation and then incubated with Biolock (IBA) for 20 minutes before StrepTactin Sepharose resin (IBA) was added. After overnight incubation at 4°C the proteins were purified on Poly-Prep chromatography columns (BioRad) according to instructions of the manufacturers of the affinity resins. Proteins were eluted from the resin using D(+)-Biotin (Roth). Proteins were analyzed for correct size and purity by sodium dodecyl sulfate polyacrylamide gel electrophoresis (SDS-PAGE) on a 12% gel and under reducing conditions, stained with GelCode Blue Stain Reagent (Thermo Scientific).

### Nanoparticle expression, purification, and NA-NP assembly

Mi3 and lumazine synthase were expressed in BL21 cells (Novagen) from the pET28a-C-tag-SpyCatcher-Mi3 (Addgene #112255) and pET15b-StrepTag-SpyCatcher-LS (Novagen) plasmids. Cells transformed with these plasmids were grown at 37°C in LB medium until they reached log phase (OD_600_∼0.8) and then expression was induced by adding isopropyl-β-d-thiogalactopyranoside (IPTG; GIBCO BRL) to a final concentration of 0.5 mM. Cultures were incubated for 16 hours at 22°C, then pelleted and resuspended in lysis buffer (50mM HEPES, 150mM NaCl, 0.1% Triton X-100, 0.1 mg/mL Lysozyme, cOmplete Protease Inhibitor (Roche)). Samples were then sonicated on ice in four rounds of 30 seconds. Debris was removed by ultracentrifugation and supernatant was incubated overnight with CaptureSelect C-Tag affinity matrix (Thermo Scientific; Mi3 purification) or StrepTactin Sepharose resin (IBA; LS purification). Nanoparticles were purified according to manufacturer’s instructions. Proteins were analyzed on 12% SDS-PAGE gel with GelCode Blue (Thermo Scientific). NA-NP nanoparticles were assembled by co-incubation of SpyTag-NA and SpyCatcher-NP in 50 mM HEPES and 150 mM NaCl pH 7.4 For 16 hours at room temperature. Conjugation of NA to the nanoparticles was confirmed with SDS-PAGE on a 12% gel under reducing conditions, stained with GelCode Blue (Thermo Scientific).

### Generation of mAbs

Harbour H2L2 transgenic mice (www.harbourbiomed.com) were immunized with six doses of 25 µg purified recombinant NA proteins per mouse in two-week intervals. One group received a mix of unconjugated N1 WI13 and N2 GE18 proteins and the other group received the N1 and N2 proteins conjugated to nanoparticles. For NA-NP vaccination we alternated between NA-Mi3 and NA-LS immunizations. The first immunization was prepared with Stimune Adjuvant (Prionics) according to manufacturer’s instructions and the booster immunizations were prepared with Ribi (Sigma) adjuvant. Immunizations were administered subcutaneously in the left and right groin (50 µl each) and intraperitoneally (100 µl). Four days after the last injection, spleen and lymph nodes were harvested and hybridomas were generated as described previously.^53^ Hybridoma supernatants were screened for reactivity against the NA antigens and negatively selected for reactivity against RSV G containing the same tetramerization domain and tags in ELISA. Antibodies from the hybridomas selected for further development were subcloned and produced in small scale (as described^54^) for further analysis in ELISA, MUNANA, and infection neutralization assays. The mouse immunization experiments were done under the animal permit 2010806, approved by the Dutch central committee for animal experiments.

### Production of human monoclonal antibodies

For recombinant mAb production, the genes encoding the heavy and light chain variable regions were sequenced (sequences are available on request). Human codon-optimized cDNA (Genscript, USA) encoding these regions was cloned into pCAGGS expression plasmids containing the human IgG1 heavy chain and Ig kappa light chain constant regions.^55^ Recombinant hIgG1 12F5, 21H8, 45D9, 1G01^19^, and isotype control (anti-RSV-F mAb Palivizumab^56^) were produced in HEK293 FreeStyle 293-F cells (Gibco) following transfection with pairs of the IgG1 heavy and light chain expression plasmids similar as described above for production of recombinant NA proteins. Antibodies were purified from the cell culture supernatants using Protein-A resin (GE Healthcare) according to manufacturer’s instructions. The antibodies were eluted using 0.1 M citric acid at pH 3.0 and neutralized using 1 M Tris-HCl at pH 8.8.

### Viruses

Mouse-adapted A/Belgium/1/2009 (H1N1; referred to as Bel/09) and A/Victoria/3/75 (H3N2 in the genetic background of PR8; referred to as X47) virus strains were used in mouse challenge experiments. These virus strains were amplified on Madin-Darby canine kidney (MDCK) cells in serum-free Dulbecco’s Modified Eagle medium (DMEM) supplemented with non-essential amino acids, 2 mM L-glutamine and 0.4 mM sodium pyruvate in the presence of 2 μg/mL TPCK-treated trypsin (Sigma) at 37 °C in 5% CO2. Ninety-six hours after virus inoculation, the culture medium was collected, cell debris was removed by centrifugation for 10 min at 2500 rcf at 4°C, and the virus was pelleted from the supernatants by overnight centrifugation at 30,000 rcf at 4°C. The pellet was resuspended in cold sterile 20% glycerol in PBS, aliquoted and stored at −80°C until used.

A/Netherlands/602/2009 (H1N1; referred to as H1N1pdm09), A/Perth/16/2009 (H3N2; referred to as PE09), A/Bilthoven/1761/1976 (H3N2 in the genetic background of A/Puerto Rico/8/34/Mount Sinai; referred to as VI75) were used in *in vitro* assays. The virus strains were amplified on Madin-Darby canine kidney (MDCK) cells in Opti-MEM reduced serum medium in the presence of 1 μg/mL TPCK-treated trypsin (Sigma) at 37 °C in 5% CO2. 48-72 hours after virus inoculation, the cell culture medium was collected, cell debris was removed by centrifugation for 10 min at 2500 rcf at 4°C. Supernatants containing the virus were aliquoted and stored at −80°C until used. For use in ELISA or NK cell activation assays, the virus stocks were further purified by incubation with Capto Core 700 (Cytiva) and concentrated using centrifugal filter units (Amicon) following manufacturer’s instructions.

### Enzyme-Linked Immunosorbent Assay

Antibody binding to NA and RSV G was determined by Enzyme-Linked Immunosorbent Assay (ELISA). NUNC MaxiSorp plates (Thermo Scientific) were coated with recombinant proteins or virus stocks diluted in PBS with calcium and magnesium, and incubated overnight at 4°C. The plates were washed three times with PBS + 0.05% Tween-20 and blocked for one hour with 3% bovine serum albumin and 0,1% Tween-20 in PBS. Serum samples, hybridoma supernatants, or purified recombinant mAbs were serially diluted in blocking buffer, added to the plates, and incubated for two hours at room temperature. After incubation, the plates were washed four times and then incubated for 1 hour with secondary antibodies diluted in blocking buffer; for serum samples and hybridoma supernatants a mix of HRP-conjugated anti-rat antibodies (IgG1; IgG2B; IgG2C) (*Absea Biotechnology*) was used, and for human hIgG1 we used rabbit-anti-human-HRP (Dako). To control for the antigen coating levels, separate wells were incubated with StrepMAB-classic HRP Conjugate (IBA) for one hour. Plates were then washed three times and incubated with TMB (BioFX) for 3-5 minutes before stopping the reaction with H_2_SO_4_. Read-out was performed using BioSPX 800 TS Microplate reader (BioTek) at 450 nm. Where indicated, values were corrected for the antigen coating levels as determined based on the StrepMAB values.

### NA peptide array analysis

We determined the linear epitopes of the mAbs with an ELISA-based peptide array analysis. The peptide arrays were derived from A/New York/18/2009 (H1N1) (NR-19249; *BEI Resources*) and A/Perth/16/2009 (H3N2) (NR-19267; *BEI Resources*) and contained 13-to 15-meric peptides with 11 amino acids overlap spanning the whole NA sequences. We combined peptides into vertical and horizontal pools (Table S1) and coated them in MaxiSorp plates (Thermo Scientific) to perform ELISAs with the mAbs similar to described above. From the combined reactivity in the vertical and horizontal pools we delineated the reactive peptides and confirmed the results by ELISA assays against the single peptides using a dilution range of the mAbs.

### MUNANA NA inhibition assay

To measure the ability of the mAbs to inhibit NA activity, we used a MUNANA assay, in which NA hydrolyzes the substrate 2ʹ-(4-methylumbelliferyl)-α-d-N-acetylneuraminic acid (MUNANA; Sigma-Aldrich) to the fluorescent 4-methylumbelliferone (4-MU).^57^ Since it uses a small molecule substrate, inhibition is only observed with antibodies that directly target the NA catalytic site. The mAbs were serially diluted in NA reaction buffer (50 mM Tris-HCl, 4 mM CaCl2, pH 6.0) in a flat-bottom 96-well black plate (Greiner Bio-One). An equal volume of H1N1 or H3N2 virus (corresponding to 80% maximum activity in MUNANA assay) diluted in the same buffer was added to each well and the plates were incubated for 30 min at RT. After incubation of virus with the mAbs, reaction buffer containing 200 µM MUNANA was added to each well and the plate was incubated at 37°C for one hour. The reaction was terminated by adding the stop solution (0.1 M glycine, 25% ethanol, 0,1% Triton-X100 pH 10.7). Fluorescence was measured immediately after stopping the reaction using Promega GloMax Explorer with excitation filter 365 nm and emission filter 415-445 nm.

### NA inhibition in ELLA

The Enzyme-Linked Lectin Assay (ELLA) was used to measure the ability of the mAbs to inhibit NA sialidase activity against a large substrate. Fetuin (Sigma) was coated in wells of MaxiSorp plates (Thermo Scientific) and incubated overnight at 4°C. The plates were then washed three times with PBS containing 0.05% Tween-20 (PBS-T) and blocked with 3% BSA for 1 hour. MAbs were serially diluted in NA reaction buffer (50 mM Tris-HCl, 4 mM CaCl2, pH 6.0) and mixed with an equal volume of the H1N1 or H3N2 IAV viruses (diluted to concentrations corresponding to 80% maximum activity in ELLA assay). The mAb/virus mixture was incubated for 30 min at RT, transferred to the fetuin-coated plates, and incubated for 4 hours at 37°C. The plates were then washed three times with PBS-T and incubated for 1 h with PNA-biotin (1:750 in PBS + 1% BSA). The plates were again washed three times with PBS-T and incubated for 1 h with streptavidin-HRP (1:1000 in PBS + 1% BSA) After three washes with PBS-T, TMB substrate was added, and the plates were incubated for 3-5 min before the reaction was stopped by the addition of H_2_SO_4_. Absorbance was measured using BioSPX 800 TS Microplate reader (BioTek) at 450 nm.

### Virus neutralization assays

Virus neutralization was assessed using a luciferase reporter assay. HeLa R19 cells were seeded in 96 well plates (10^4^ cells/well). Cells were transfected the next day with the pHH-Gluc luciferase reporter plasmid^58^, using FuGENE transfection reagent (Promega) according to manufacturer’s instructions. The pHH-Gluc plasmid contains the gaussian luciferase gene, flanked by 3’ and 5’ untranslated regions of the IAV NP genome segment, under control of the RNA polymerase I promoter in negative sense orientation. Generation of luciferase-encoding mRNAs and thus of luciferase protein is only observed after infection of transected cells with IAV. After a further 24 hour incubation at 37°C and 5% CO2, mAbs were serially diluted in Opti-MEM in a 96 well plate. H1N1 or H3N2 virus was diluted to a concentration corresponding to 90% maximum signal and added to the mAbs. Virus-mAb mixture was incubated for 15 min and meanwhile the luciferase-transfected cells were washed with PBS. The virus-mAb mixture was transferred to the cells and incubated for 16 hours at 37°C and 5% CO_2_.

### NK cell activation assay

To assess the ability of the mAbs to induce NK cell activation, we coated plates with NA (recombinant protein or virus), allowed the mAbs to bind the antigens, then incubated the NK cells and measured CD107a in flow cytometry as a measure of activation.^59^ Recombinant soluble NA proteins or purified virus stocks were diluted in PBS with calcium and magnesium and incubated in MaxiSorp plates (Thermo Scientific) overnight at 4°C. The plates were washed with PBS and blocked for 30 minutes at 37°C with 5% BSA in PBS. MAbs were serially diluted in PBS with 1% BSA, added to the plates, and incubated for 1h at RT. NK92 cells were diluted to 10^6^ cells/ml in OptiMEM reduced serum medium (Gibco) supplemented with 1x Protein Transport Inhibitor Cocktail (Invitrogen) and V450 mouse anti-human CD107a (BD Biosciences; 1:400). Plates were washed with PBS and 100 µl of the cell suspension was added per well. Cells were incubated in the plates for 5 hours at 37°C with 5% CO_2_. After 5 hours, the cells were washed and stained for flow cytometry with PE mouse anti-human CD56 (BD Biosciences; 1:100) and LIVE/DEAD Fixable Near IR (876) stain (Invitrogen; 1:1000) diluted in FACS buffer (PBS with 0.05% BSA and 0.02% NaN_3_). The cells were incubated at 4°C for 45 min, then washed, fixed with 3% paraformaldehyde for 10 min, and resuspended in FACS buffer. Measurements were performed on a BC CytoFLEX LX (BD Biosciences) and results were analyzed using FlowJo. Events were gated on single, viable and CD56 positive cells, and the percentage of CD107a positive cells was determined within this population.

### Complement deposition assay

The ability of the mAbs to induce complement activation was measured as the deposition of the C3b complement protein on NA-coated microspheres.^60^ Biotinylated NA proteins were conjugated to neutravidin-labelled FluoSpheres 580/605 (Invitrogen) in a 1:1 ratio (µg antigen:µl microspheres) by overnight incubation at 4°C while shaking. The microspheres were washed with 2% BSA in PBS and resuspended in PBS with 0.1% BSA. 10^6^ microspheres were added per well to a 96-well plate containing serial dilutions of the mAbs and incubated for 2h at 37°C to form immune complexes. The immune complexes were then either incubated with anti-human IgG Alexa Fluor 647 (Invitrogen; 1:400, incubated for 1h at 4°C) to determine the antibody binding levels or guinea pig complement (Sigma; 1:25, incubated for 15 min at 37°C) followed by washing in FACS buffer and labelling with FITC goat anti-guinea pig complement C3 (MP Biomedicals; 1:100, incubated for 30 min at 4°C). The microspheres were washed twice with PBS, fixed with 3% paraformaldehyde for 15 min at RT, washed again, and resuspended in FACS buffer. Acquisition was performed on a BC CytoFLEX LX (BD Biosciences). Events were gated on single microspheres and the median fluorescence intensity of the FITC (C3 binding) or Alexa Fluor 647 (antibody binding) signals was determined as a final readout.

### Phagocytosis assay

Phagocytosis induced by mAb binding was assessed by determining phagocytic uptake of microsphere-based immune complexes by THP-1 cells, a human monocyte cell line. Immune complexes were prepared as described above for the complement deposition assay. Then, 5x10^4^ THP-1 cells in RPMI-1640 (ADCC) were added to each well and the cells were incubated with the immune complexes for 1h at 37°C and 5% CO_2_. All plates were washed twice with PBS and fixed with 3% paraformaldehyde for 15 min at RT. Then, the beads were washed and resuspended in FACS buffer. Acquisition was performed on a BC CytoFLEX LX (BD Biosciences). Events were gated for single cells and cells containing the fluorescent microspheres (% beads^+^ cells). The median fluorescence intensity of the beads in the beads positive cell population was determined (gMFI (beads^+^ cells)). A phagoscore was calculated with the following equation:

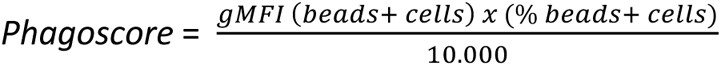

### In vivo challenge studies

Six-to 8-week-old female BALB/c mice (Charles River) were housed under specific-pathogen-free conditions with food and water *ad libitum*. One day prior to virus challenge, mice were treated with mAbs by intranasal administration. The day after, mice were weighed to obtain baseline measurements, before being challenged with a 2LD_50_ or 4LD_50_ dose of mouse-adapted Bel/09 or X47 virus. Influenza virus inoculations were performed under sedation with a mixture of ketamine (Eurovet, Netamik, 10 mg/kg) and xylazine (Bayer, Rompun, 60 mg/kg) administered intraperitoneal, and a total of 50 μl was instilled equally into the nostrils of the mouse. After infection, the body weight of the mice was determined daily for two weeks. Animals that had lost ≥25% of their original body weight were humanely euthanized by cervical dislocation. The mouse challenge experiments were conducted according to the Belgian legislation (Belgian Law 14/08/1986 and Belgium Royal Decree 06/04/2010) and European legislation on protection of animals used for scientific purposes (EU directives 2010/63/EU and 86/609/EEC). Experimental protocols were all approved by the Ethics Committee of the Vlaams Instituut voor Biotechnologie (VIB), Ghent University, Faculty of Science (permit numbers EC2021-032 and EC2022-104).

### Statistical analysis

Relative body weights, expressed as a percentage of baseline body weight on the day of infection, were calculated for each mouse at various time points post-infection (days 1-14; dpi). These data were presented using GraphPad version 8.00, with a mean (n=6) and error bar (s.e.m) displayed for each group at each time point.

Relative body weights were analyzed as repeated measurements using the method of residual maximum likelihood (REML) as implemented in Genstat v22 (VSN International). Briefly, a linear mixed model with treatment, time and the treatment × time interaction as fixed terms, and the mouse × time interaction term set as random term, was fitted to data. The term mouse × time represents the residual error term with dependent errors because the repeated measurements are taken of the same mouse, causing correlations among observations. The autoregressive correlation model of order 1 (AR1) with times of measurement set at equal intervals, was selected as the best-fitted model based on Akaike’s information criterion coefficient. The AR1 covariance model assumes that the correlation between observations decays as the measurements are collected further apart in time. The first order refers to the fact that recent (t-1) observations affect the current observations made at timepoint t. Additional options selected to get a best-fitting model included an allowance of unequal variances across time. The significance of the fixed terms in the model and the significance of changes in the difference between treatment effects over time were assessed using an approximate F-test as implemented in Genstat v22 (VSN International).

Mice that had lost ≥25% body weight were considered as death events at the measuring time point. Survival data were fit into the survival table format in GraphPad version 8.00. Differences in survival were determined using the Mantel-Cox log-rank test.

### Immunofluorescence imaging

To determine binding of the mAbs to full-length, membrane-bound NA, we infected MDCK cells, stained with the mAbs, and imaged using confocal microscopy. MDCK cells were seeded in 18-well glass-bottom chambers (Ibidi) and incubated at 37°C and 5% CO_2_. After 24 hours the cells were infected with H1N1pdm09 or H3N2 VI75 viruses in OptiMEM. Cells were incubated for 8 hours at 37°C and 5% CO_2_, then washed with PBS and fixed with 4% paraformaldehyde in PBS for 20 minutes at RT. Cells were then washed three times with PBS containing 10 mM glycine, blocked with 2% BSA in PBS for 30 minutes, and washed again with PBS containing 10 mM glycine. Cells were stained with the NA mAbs at 10 µg/ml, followed by goat anti-human IgG AlexaFluor 488 (Invitrogen), both for 1 hour at RT. Then, cells were stained with anti-influenza nucleoprotein monoclonal antibody (HB65) harvested from hybridoma H16-L10-4R5 (ATCC), followed by donkey anti-mouse IgG Alexa Fluor 647 (Invitrogen). Nuclei were labelled with DAPI (Invitrogen). Cells were washed in between staining steps and after the last incubation and imaging buffer was added (2x SSC buffer with 10mM Tris (pH = 8), 0,4% glucose (w/v) supplemented with glucose oxidase (Sigma-Aldrich) and catalase (Sigma-Aldrich)). For intracellular staining of NA, cells were permeabilized by incubation with 0.2% Triton X-100 in PBS for 5 minutes prior to the incubation with the NA mAbs. For surface staining of NA, the permeabilization was performed right before addition of the anti-nucleoprotein mAb. Imaging was performed using the Olympus SpinSR spinning disc system with SoRa disk and a 60x oil immersion 1.5NA objective with additional 3.2x magnification. Images were analyzed using Fiji ImageJ.

