## Supplementary figures and tables for "Fc-dependent protective efficacy of non-inhibitory antibodies targeting influenza A virus neuraminidase is limited by epitope availability"

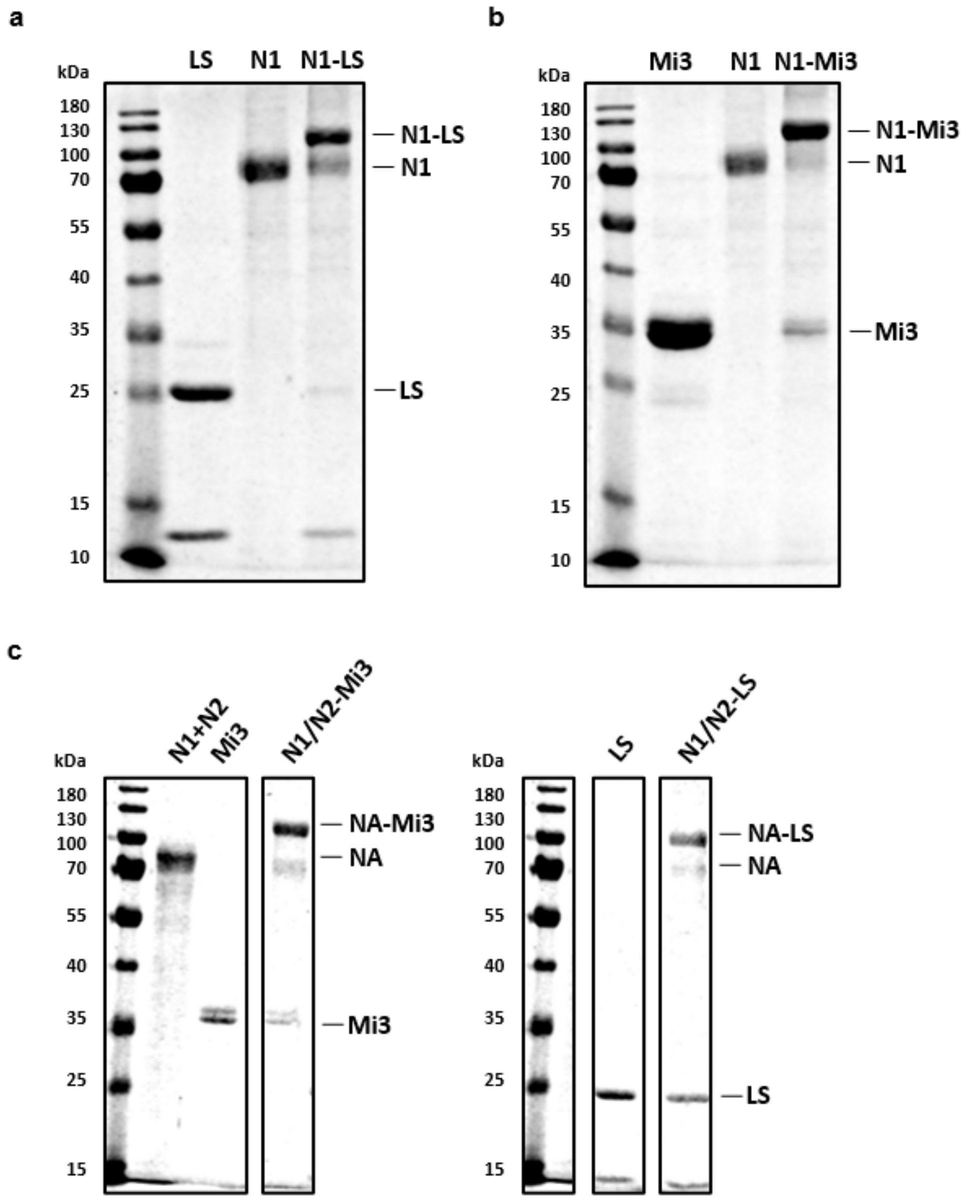


**Supplementary figure 1. Confirmation of NA-NP conjugation.** Conjugation of N1 to LS (a) and Mi3 (b) nanoparticles demonstrated by reduced electrophoretic mobility on SDS-PAGE. (c) Conjugation of both N1 and N2 NAs to Mi3 (left) and LS (right) for immunizations.

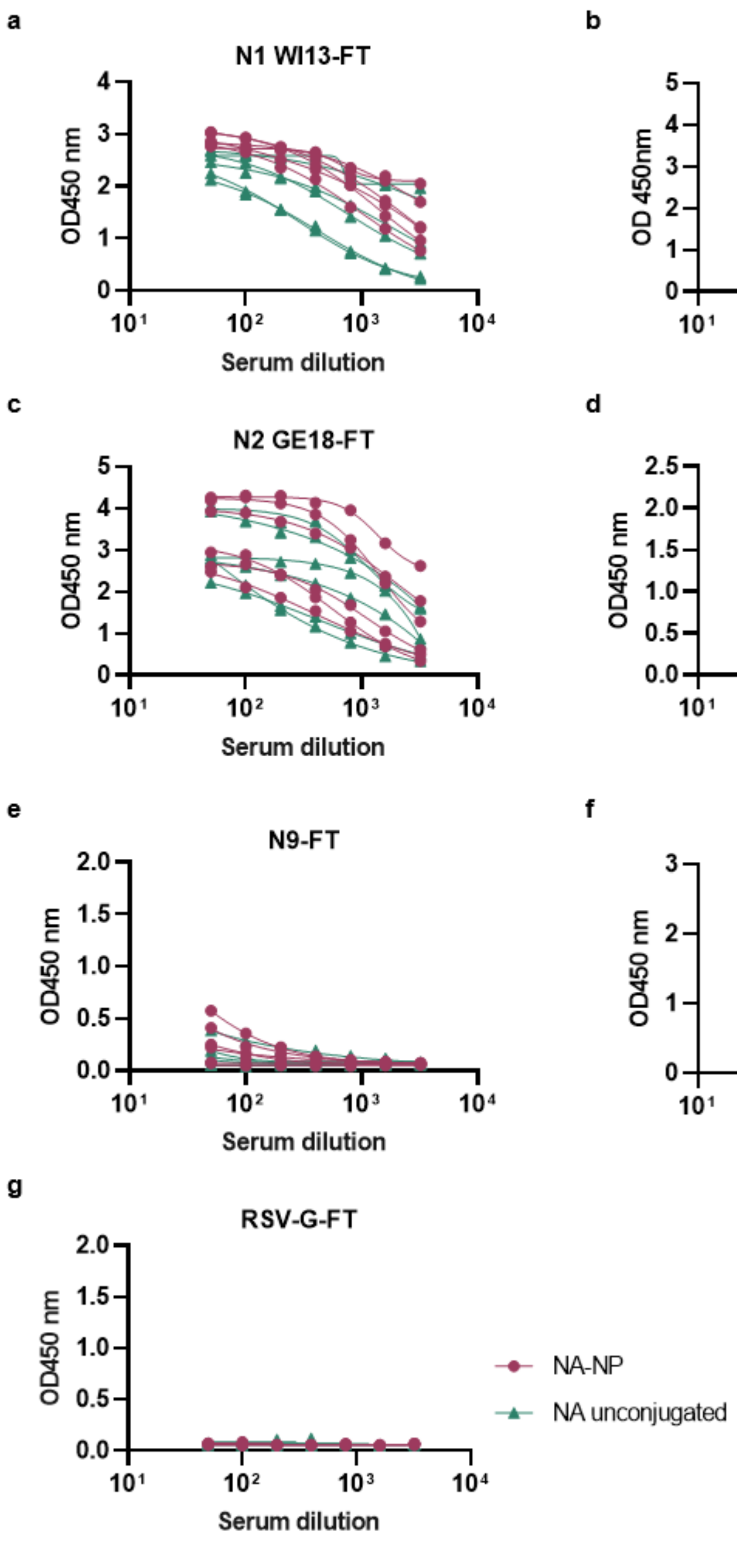


**Supplementary figure 2. NA(-NP) immunizations induce high serum reactivity against NAs.** H2L2 mice were immunized with unconjugated NA or NA-NP. Reactivity against different NA subtypes (a-e) and control antigens (f and g) was quantified in ELISA using serial dilutions of sera collected after four immunizations. NA antigens used in ELISA contained a different tetramerization domain and purification tag from the antigens used for immunization (indicated with FT) to which the mice are naive, as to measure the reactivity against the NA only. RSV-G-SPY (f) contains the tetramerization domain, purification tag, and spy tag present in the recombinant NA proteins used in the immunizations.

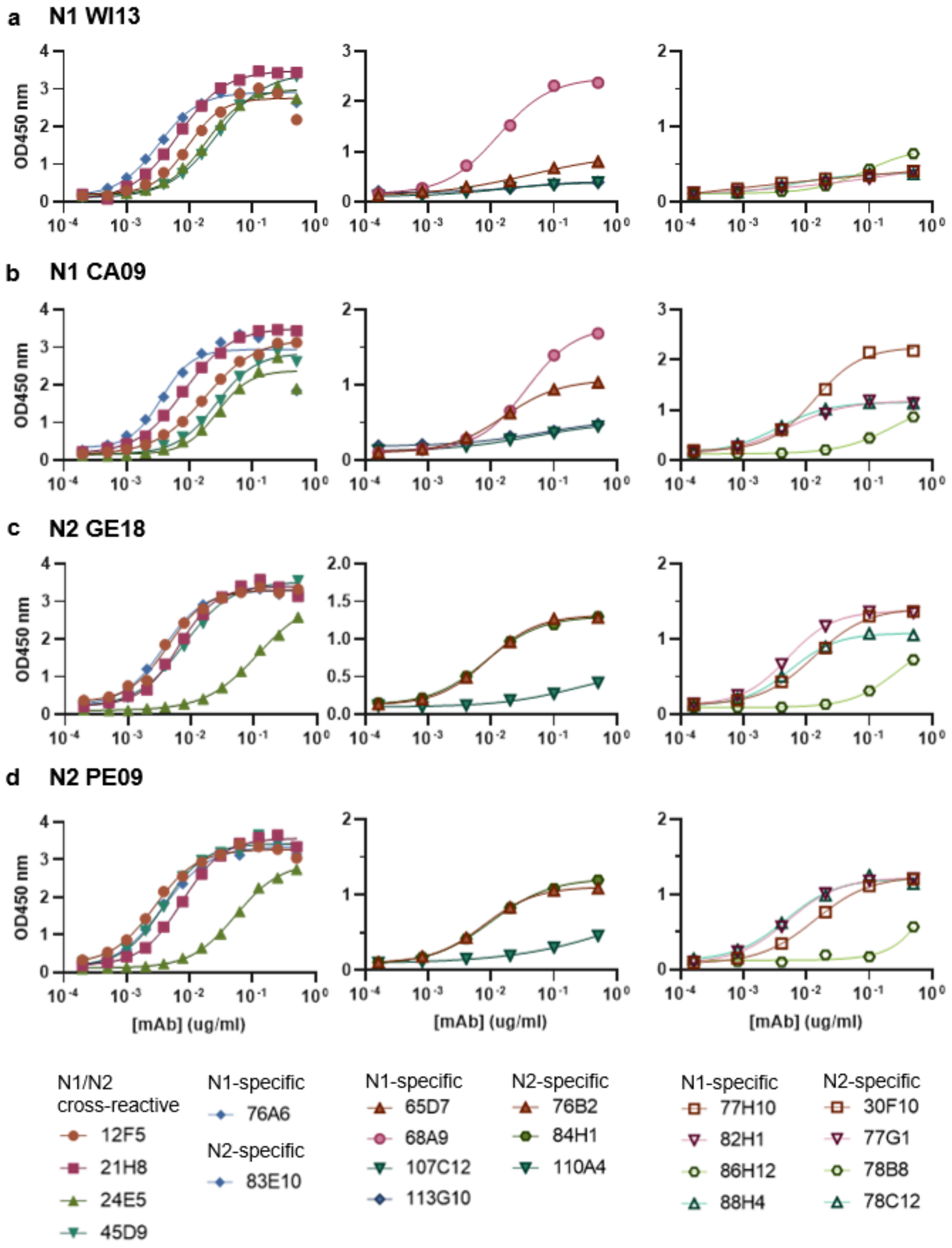


**Supplementary figure 3. Quantification of binding of the rIgG mAbs against NA in ELISA.** Binding of the mAbs isolated from hybridoma supernatants was measured in ELISA against N1 WI13 (a), N1 CA09 (b), N2 GE18 (c), and N2 PE09 (d) using serial dilutions of mAbs (preliminary data based on a single experiment with no replicates).

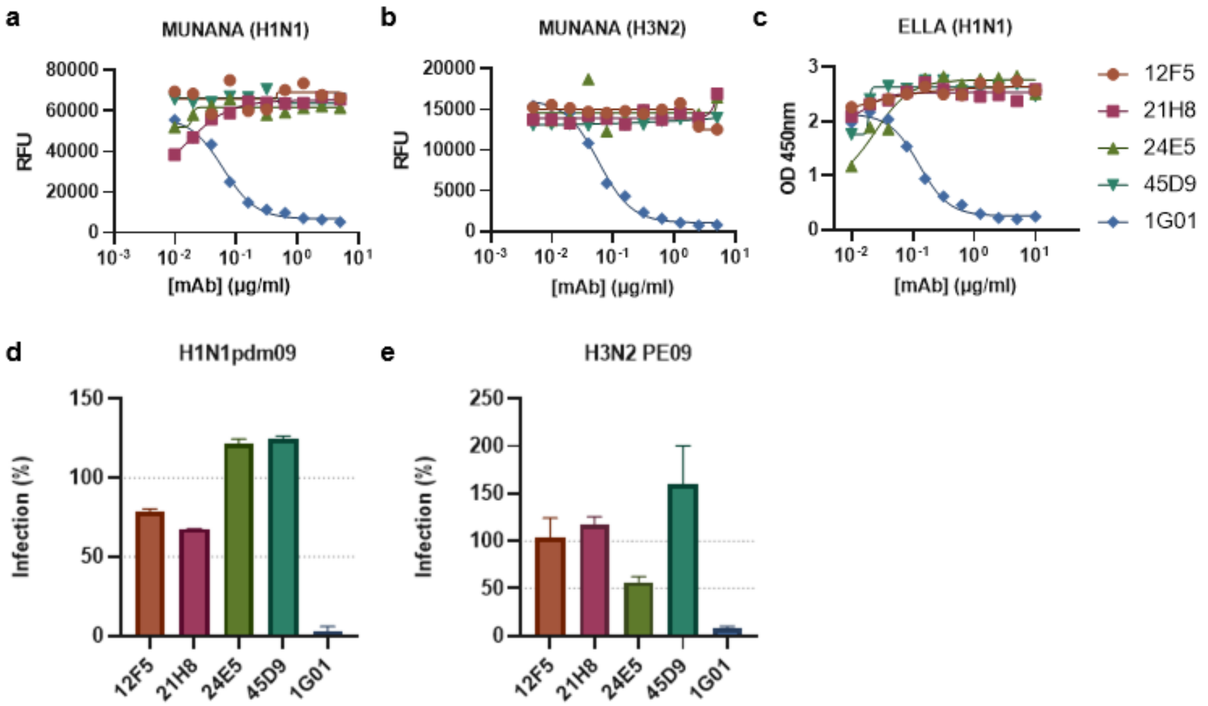


**Supplementary figure 4. Cross-reactive rat IgG mAbs do not inhibit NA activity or infection *in vitro*.** Functional characterization of the IgG mAbs with rat Fc domains and positive control hIgG1 1G01 in NA inhibition assays and infection inhibition assay. Inhibition of NA activity was measured in MUNANA assay with H1N1pdm09 (a) and H3N2 VI75 (b) and in ELLA with H1N1pdm09 (c) using serial dilutions of the mAbs (preliminary data based on a single measurement with no replicates). Inhibition of infection was determined in a single-round infection assay with luciferase reporter read-out with H1N1pdm09 (d) and H3N2 PE09 (e) normalized to mock-treated controls (representative data of two independent assays; mean ±SD of three replicate samples).

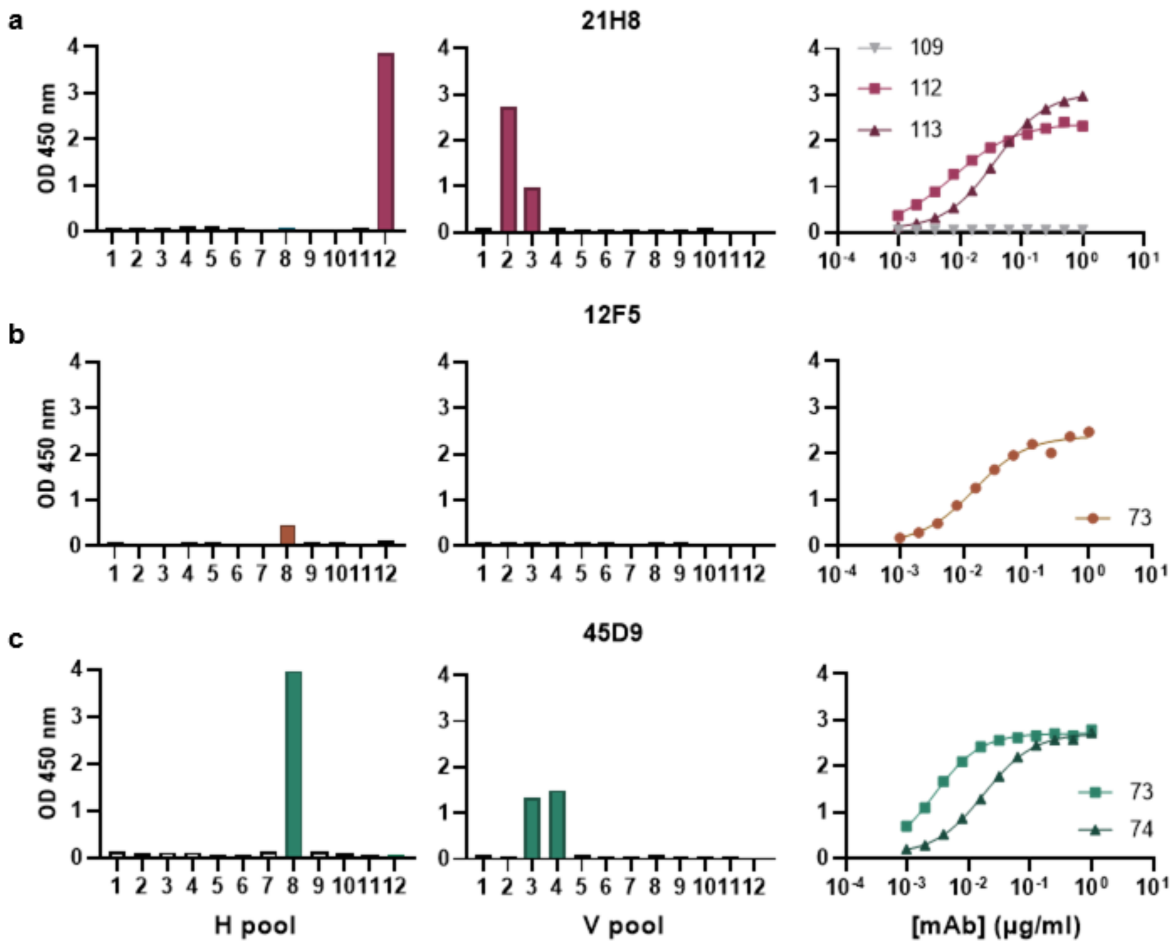


**Supplementary figure 5. Peptide array to determine binding epitopes of the cross-reactive mAbs in N1.** Peptide array derived from A/New York/18/2009 (H1N1). Reactivity of the mAbs (a, 21H8; b, 12F5; c, 45D9) against pooled peptides was measured in ELISA (left and middle graphs) and confirmed for the selected peptides (right). (d) Allocation of peptides in horizontal pools (H1-H12) and vertical pools (V1-V10) is shown in table 1.

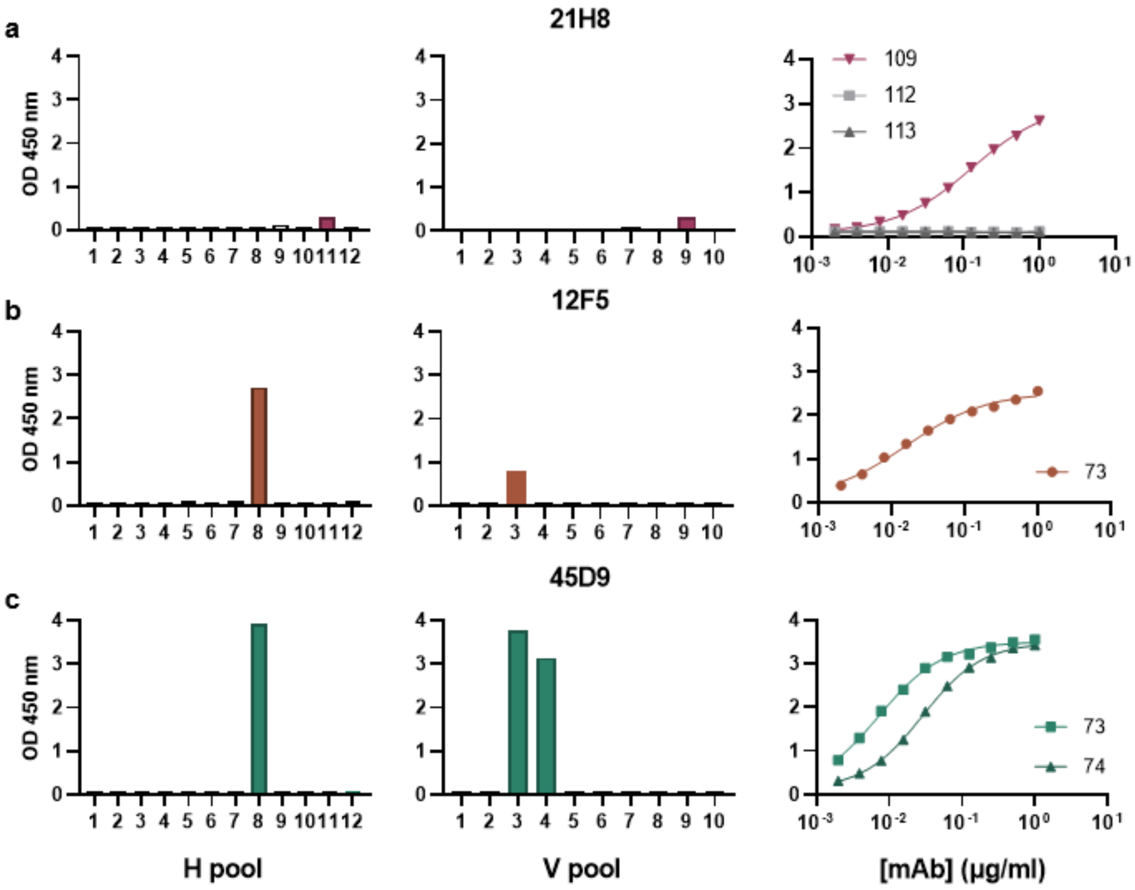


**Supplementary figure 6. Peptide array to determine binding epitopes of the cross-reactive mAbs in N2.** Peptide array derived from A/Perth/16/2009 (H3N2). Reactivity of the mAbs (a, 21H8; b, 12F5; c, 45D9) against pooled peptides was measured in ELISA (left and middle graphs) and confirmed for the selected peptides (right). Allocation of peptides in horizontal pools (H1-H12) and vertical pools (V1-V10) is shown in table 1.

```
CLUSTAL O(1.2.4) multiple sequence alignment

N2_BE92      GTCTVVMTDGSASERADTKILFIEEGKIVHISPLSGSAQHVEECSCYPRYPGVRCVCRDN    294
N2_GE18      GTCTVVMTDGNATGKADTKILFIEEGKIIHTSKLSGSAQHVEECSCYPRYPGVRCVCRDN    294
N2_FU02      GTCTVVMTDGSASGKADTKILFIEEGKIVHTSTLSGSAQHVEECSCYPRYPGVRCVCRDN    294
N2_PE09      GTCTVVMTDGSASGKADTKILFIEEGKIVHTSTLSGSAQHVEECSCYPRYPGVRCVCRDN    294
N2_HK68      GTCTVVMTDGSASGRADTRILFIEEGKIVHISPLSGSAQHVEECSCYPRYPGVRCICRDN    294
N2_VI75      GTCTVVMTDGSASGRADTKILFIEEGKIVHTSPLSGSAQHVEECSCYPRYPGVRCICRDN    294
N1_PR8       GSCFTIMTDGPSDGLASYKIFKIEKGKVTKSIELNAPNSHYEECSCYPDTGKVMCVCRDN    280
N1_KY07      GSCFTIMTDGPSNGAASYKIFKIEKGKVTKSIELNAPNFHYEECSCYPDTGTVMCVCRDN    295
N1_HU02      GSCFTVMTDGPSNGQASYKIFKMEKGKVVKSVELDAPNYHYEECSCYPDAGEITCVCRDN    275
N1_CA09      GSCFTVMTDGPSNGQASYKIFRIEKGKIVKSVEMNAPNYHYEECSCYPDSSEITCVCRDN    295
N1_WI13      GSCFTIMTDGPSDGQASYKIFRIEKGKIVKSVEMNAPNYHYEECSCYPDSSEITCVCRDN    295
             *:* .:**** :   *. :*: :*:**: :   :..   * *******    : *:****
             45D9 N2 epitope

N2_BE92      WKGSNRPIVDINVKDYSIVSSYVCSGLVGDTPRKNDSSSSSYCRNPNNEKGSHGVKGWAF    354
N2_GE18      WKGSNRPIIDINIKDHSIVSSYVCSGLVGDTPRKSDSSSSSHCLNPNNEEGGHGVKGWAF    354
N2_FU02      WKGSNRPIVDINIKDYSIVSSYVCSGLVGDTPRKNDSSSSSHCLGPNNEEGGHGVKGWAF    354
N2_PE09      WKGSNRPIVDINMKDHSIVSSYVCSGLVGDTPRKNDSSSSSHCFDPNNEEGGHGVKGWAF    354
N2_HK68      WKGSNRPVVDINMEDYSIDSSYVCSGLVGDTPRNDDRSSNSNCRNPNNERGNQGVKGWAF    354
N2_VI75      WKGSNRPVVDINVKDYSIDSSYVCSGLVGDTPRNNDRSSSSYCRNPNNEKGTHGVKGWAF    354
N1_PR8       WHGSNRPWVSFDQN-LDYQIGYICSGVFGDNPRPEDGTGSC---GPVYVDGANGVKGFSY    336
N1_KY07      WHGSNRPWVSFNQN-LDYQIGYICSGVFGDNPRPKDGKGSC---NPVTVDGADGVKGFSY    351
N1_HU02      WHGSNRPWVSFNQN-LEYQIGYICSGVFGDNPRPNDGTGSC---GPVSPNGAYGVKGFSF    331
N1_CA09      WHGSNRPWVSFNQN-LEYQIGYICSGIFGDNPRPNDKTGSC---GPVSSNGANGVKGFSF    351
N1_WI13      WHGSNRPWVSFNQN-LEYQIGYICSGVFGDNPRPNDKTGSC---GPVSSNGANGVKGFSF    351
             *:***** :.:: :  .   .*:***:.**.** .* ....   .*    *  ****:::
             45D9 N1 epitope

N2_BE92      DDGNDVWMGRTISEELRSGYETFKVIEGWSKPNSKLQINRQVIVDRGNRSGYSGIFS---    411
N2_GE18      DDGNDVWMGRTINETSRLGYETFKVVEGWSNPKSKLQINRQVIVDRGDRSGYSGIFS---    411
N2_FU02      DDGNDVWMGRTISEKLRSGYETFKVIEGWSNPNSKLQINRQVIVDRGNRSGYSGIFS---    411
N2_PE09      DDGNDVWMGRTISEKSRLGYETFKVIEGWSNPKSKLQINRQVIVDRGDRSGYSGIFS---    411
N2_HK68      DNGDDVWMGRTISKDLRSGYETFKVIGGWSTPNSKSQINRQVIVDSDNRSGYSGIFS---    411
N2_VI75      DDGNDVWMGRTISEDSRSGYETFKVIGGWSTPNSKLQINRQVIVDSDNRSGYSGIFS---    411
N1_PR8       RYGNGVWIGRTKSHSSRHGFEMIWDPNGWTETDSKFSVRQ-DVVAMTDWSGYSGSFVQHP    395
N1_KY07      KYGNGVWIGRTKSNRLRKGFEMIWDPNGWTDTDSDFSVKQ-DVVAITDWSGYSGSFVQHP    410
N1_HU02      KYGNGVWIGRTKSTNSRSGFEMIWDPNGWTGTDSSFSVKQ-DIVAITDWSGYSGSFVQHP    390
N1_CA09      KYGNGVWIGRTKSISSRNGFEMIWDPNGWTGTDNNFSIKQ-DIVGINEWSGYSGSFVQHP    410
N1_WI13      KYGNGVWIGRTKSISSRKGFEMIWDPNGWTGTDNKFSIKQ-DIVGINEWSGYSGSFVQHP    410
               *:.**:*** .   * *:* :    **:  ... .:.:  :*   : ***** *

N2_BE92      -VEGKSCINRCFYVELIRGRKQETEVWWTSNSIVVFCGTSGTYGTGSWPDGADINLMPI-    469
N2_GE18      -VEGKSCINRCFYVELIRGRKEETKVLWTSNSIVVFCGTSGTYGTGSWPDGADLNLMHI-    469
N2_FU02      -VEGKSCINRCFYVELIRGRKQETEVLWTSNSIVVFCGTSGTYGTGSWPDGADINLMPI-    469
N2_PE09      -VEGKSCINRCFYVELIRGSKEETEVLWTSNSIVVFCGTSGTYGTGSWPDGADINLMPI-    469
N2_HK68      -VEGKSCINRCFYVELIRGRKQETRVWWTSNSIVVFCGTSGTYGTGSWPDGANINFMPI-    469
N2_VI75      -VEGKSCINRCFYVELIRGREQETRVWWTSNSIVVFCGTSGTYGTGSWPDGADINIMPI-    469
N1_PR8       ELTGLDCMRPCFWVELIRGRPKEK-TIWTSASSISFCGVNSDTVDWSWPDGAELPFSIDK    454
N1_KY07      ELTGLDCIRPCFWVELVRGLPRENTTIWTSGSSISFCGVDSDTANWSWPDGAELPFTIDK    470
N1_HU02      ELTGLDCIRPCFWVELIRGRPKES-TIWTSGSSISFCGVNSDTVSWSWPDGAELPFTIDK    449
N1_CA09      ELTGLDCIRPCFWVELIRGRPKEN-TIWTSGSSISFCGVNSDTVGWSWPDGAELPFTIDK    469
N1_WI13      ELTGLDCIRPCFWVELIRGRPEEN-TIWTSGSSISFCGVNSDTVGWSWPDGAELPFTIDK    469
              : * .*:. **:***:**  .*. . *** * : ***...     ******:: :
                                      21H8 N2 epitope    21H8 N1 epitope
```

**Supplementary figure 7. Alignment of N1 and N2 NAs with epitopes of 45D9 and 21H8 mAbs indicated.** Epitopes determined in peptide arrays (Fig. S5 and S6). When the antibodies recognized two peptides, the overlapping sequence is indicated with a darker shade and the remaining parts of the sequences with a lighter shade.

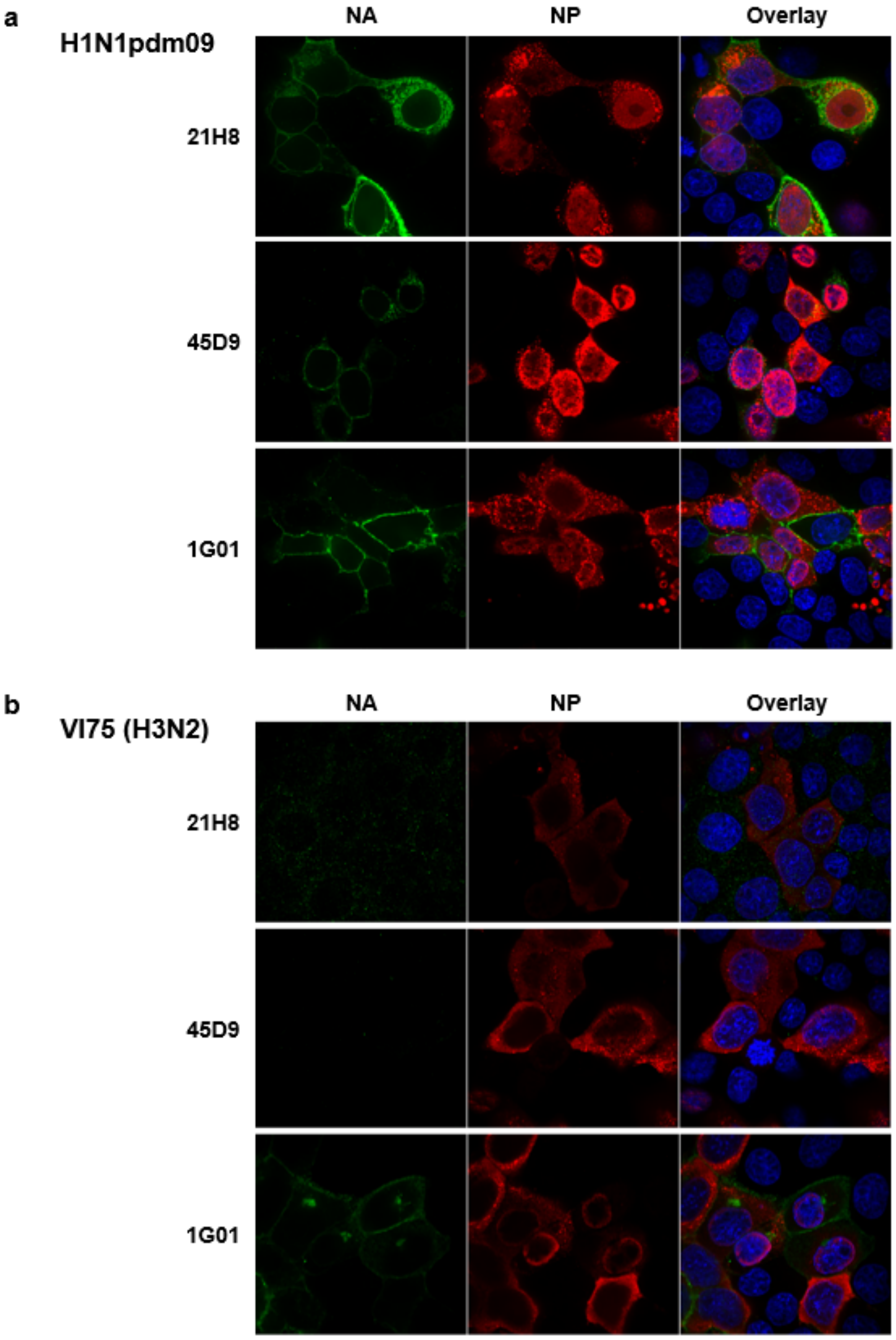


**Supplementary figure 8. Intracellular staining of infected cells with 21H8 and 45D9 mAbs.** MDCK cells were infected with H1N1pdm09 (a) and H3N2 VI75 (b). Fixed and permeabilized cells were stained intracellularly with anti-NA mAbs 21H8, 45D9, and 1G01 (green) and anti-NP mAb (red). Nuclei were labelled with DAPI (blue).

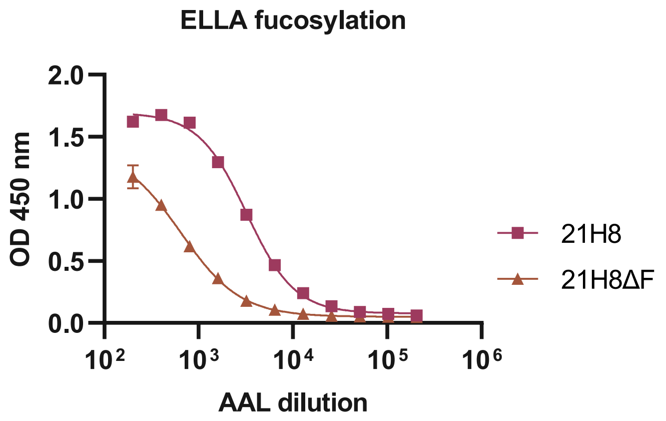


**Supplementary figure 9. RMD co-transfection reduces fucosylation of hIgG1 21H8 mAb.** HEK293-F cells were co-transfected with plasmids encoding 21H8 hIgG1 mAb and RMD to reduce core fucosylation of the Fc glycan. Fucosylation of the mAb produced with RMD co-transfection (21H8ΔF) and produced in the regular way without RMD (21H8) was quantified in an ELLA assay using AAL lectin (mean ±SD of two replicate measurements).

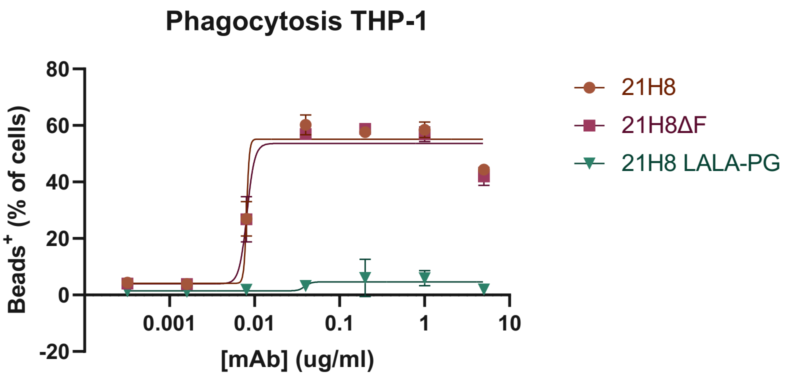


**Supplementary figure 10. Phagocytosis activation by modified 21H8 mAbs in THP-1 cells.** Phagocytosis of microsphere-based immune complexes containing N1 WI13 and the mAbs by THP-1 monocytes was measured by flow cytometry. The phagoscores were calculated based on the percentage of THP-1 cells positive for the uptake of microspheres and the gMFI of that population as detailed in the Materials and Methods section (mean ±SD of three replicate measurements).

**Supplementary table 1. Allocation of peptides from peptide arrays in horizontal and vertical pools.**

| | V1 | V2 | V3 | V4 | V5 | V6 | V7 | V8 | V9 | V10 |
|---|---|---|---|---|---|---|---|---|---|---|
| **H1** | 1 | 2 | 3 | 4 | 5 | 6 | 7 | 8 | 9 | 10 |
| **H2** | 11 | 12 | 13 | 14 | 15 | 16 | 17 | 18 | 19 | 20 |
| **H3** | 21 | 22 | 23 | 24 | 25 | 26 | 27 | 28 | 29 | 30 |
| **H4** | 31 | 32 | 33 | 34 | 35 | 36 | 37 | 38 | 39 | 40 |
| **H5** | 41 | 42 | 43 | 44 | 45 | 46 | 47 | 48 | 49 | 50 |
| **H6** | 51 | 52 | 53 | 54 | 55 | 56 | 57 | 58 | 59 | 60 |
| **H7** | 61 | 62 | 63 | 64 | 65 | 66 | 67 | 68 | 69 | 70 |
| **H8** | 71 | 72 | 73 | 74 | 75 | 76 | 77 | 78 | 79 | 80 |
| **H9** | 81 | 82 | 83 | 84 | 85 | 86 | 87 | 88 | 89 | 90 |
| **H10** | 91 | 92 | 93 | 94 | 95 | 96 | 97 | 98 | 99 | 100 |
| **H11** | 101 | 102 | 103 | 104 | 105 | 106 | 107 | 108 | 109 | 110 |
| **H12** | 111 | 112 | 113 | 114 | 115 | | | | | |

**Supplementary Table 2. Statistical analysis of the body weight differences in figures 5a and 5c.**

| | F pr. [a] | |
|---|---|---|
| **Comparison** | **Bel09** | **X47** |
| **21H8 vs 45D9** | <0.001 | 0.058 |
| **21H8 vs NC** | <0.001 | 0.040 |
| **45D9 vs NC** | <0.001 | 0.007 |
| **21H8 vs PC** | <0.001 | / |
| **45D9 vs PC** | <0.001 | / |
| **PC vs NC** | <0.001 | / |

[a] Statistical significance of the differences in relative body weight (= body weight day N/body weight day 0 x 100%, N = 1,2,…14) between the different groups was determined using a linear mixed model with repeated measurement, in which the 14-day measurement of the body weight is the repeated data for one individual.

**Supplementary Table 3. Statistical analysis of the body weight differences in figures 8a and 8b.**

| | F pr. [a] | |
|---|---|---|
| **Comparison** | **2LD50** | **4LD50** |
| **21H8 vs NC** | <0.001 | <0.001 |
| **21H8 vs 21H8-LALA-PG** | <0.001 | <0.001 |
| **21H8 vs 21H8ΔF** | 0.042 | 0.0671 |
| **21H8ΔF vs 21H8-LALA-PG** | <0.001 | <0.001 |
| **21H8ΔF vs NC** | <0.001 | <0.001 |
| **21H8-LALA-PG vs NC** | 0.043 | 0.261 |

[a] Statistical significance of the differences in relative body weight (= body weight day N/body weight day 0 x 100%, N = 1,2,…14) between the different groups was determined using a linear mixed model with repeated measurement, in which the 14-day measurement of the body weight is the repeated data for one individual.
